# TrAGEDy: Trajectory Alignment of Gene Expression Dynamics

**DOI:** 10.1101/2022.12.21.521424

**Authors:** Ross F. Laidlaw, Emma M. Briggs, Keith R. Matthews, Richard McCulloch, Thomas D. Otto

## Abstract

1

**Motivation:** Single-cell transcriptomics sequencing is used to compare different biological processes. However, often, those processes are asymmetric which are difficult to integrate. Current approaches often rely on integrating samples from each condition before either cluster-based comparisons or analysis of an inferred shared trajectory.

**Results:** We present Trajectory Alignment of Gene Expression Dynamics (TrAGEDy), which allows the alignment of independent trajectories to avoid the need for error-prone integration steps. Across simulated datasets, TrAGEDy returns the correct underlying alignment of the datasets, outperforming current tools which fail to capture the complexity of asymmetric alignments. When applied to real datasets, TrAGEDy captures more biologically relevant genes and processes, which other differential expression methods fail to detect when looking at the developments of T cells and the bloodstream forms of *Trypanosoma brucei* when affected by genetic knockouts.

**Availability and Implementation:** TrAGEDy is freely available at https://github.com/No2Ross/TrAGEDy, and implemented in R.

## 2 Introduction

First described in 2014 (Trapnell et al. 2014), Trajectory Inference (TI) methods order cells based on the gradual change in transcript levels in underlying biological processes captured by single cell RNA-sequencing (scRNA-seq). Cells are assigned a “pseudotime” value based on their position in the inferred trajectory, allowing differential gene expression tests over pseudotime. Various methods have been developed to identify differentially expressed (DE) genes in a biological process of interest, in some cases allowing targeted comparisons. Notably, TradeSeq (Van den Berge et al. 2020) uses negative binomial general additive models (GAM) and Wald tests to assess whether a gene is differentially expressed over pseudotime within a trajectory or between lineages of the same trajectory. Alternatively, Monocle’s BEAM (X. Qiu et al. 2017) identifies genes associated with a particular branch of a trajectory, whereas pseudotimeDE (Song and Li 2021) uses permutation to account for uncertainty in pseudotime and a zero-inflated negative binominal GAM to account for expression value dropout. By applying these techniques, dynamic gene expression patterns associated with a variety of interesting biological events have been investigated, such as regulators of myogenesis (Trapnell et al. 2014), effector gradients in CD4+ T cells (Cano-Gamez et al. 2020), and life cycle transitions of the pathogen *Trypanosoma brucei* (Briggs et al. 2021).

Some of these methods can be extended to identify DE genes between two conditions where the common biological process under analysis varies, such as before and after mutation or in an overlapping response to different stimuli (Van den Berge et al. 2020). One approach is to integrate the separate datasets together and complete a cluster-based comparison between conditions, treating cells as being at a discrete stage in development captured in each cluster. A limitation of this approach, however, is that development may be continuous, with some cells falling between stage boundaries. Another approach is to integrate the datasets together, perform TI to capture the shared, possibly branching, trajectory and differential expression tests across this pseudotime axis for each condition, using tools like TradeSeq’s conditionTest. However, the integration may force similarity between the trajectories, thus obscuring DE genes.

Alternatively, trajectory alignment can align cells from independently generated trajectories to find a common pseudotime axis that retains the original ordering of cells without dataset integration. Analysing aligned trajectories can reveal DE genes across the captured process, as well as differences between conditions. The discrete point at which the genes are DE between conditions can also be found. The first method described to perform such trajectory alignment was cellAlign (Alpert et al. 2018), which uses dynamic time warping (DTW) to align two trajectories together. Interpolated points are created across the trajectory, and gene expression values are assigned to the interpolated points - based on the cells that are in pseudotime vicinity to the points. Similarity between the conditions is assessed by computing the distance or correlation between the scaled gene expression values for the interpolated points on opposing conditions and, from this, the minimum cost path through the interpolated points is calculated using DTW.

The use of DTW imposes some constraints on the type of alignment that can be analysed. Each trajectory being compared must have the same start and end points, and each part of a trajectory must be matched to at least one point in the other trajectory. The practical implication of such limitations is that cell types that may not be represented on the opposing trajectory are nevertheless matched to a point on the other trajectory, making interpretation of the subsequent alignment difficult.

Here, we build on the work of cellAlign with Trajectory Alignment of Gene Expression Dynamics (TrAGEDy), where we make post-hoc changes to the alignment, allowing us to overcome the limitations of DTW and better reflect differences that may occur across the alignment. We also implement an approach to identify DE genes across the alignment, in order to better identify differences between the two conditions. We test TrAGEDy with a variety of simulated scRNA-seq datasets with different underlying alignments, as well datasets of *Trypanosoma brucei* lifecycle and T cell development under different genetic conditions, revealing gene expression changes undetected by current methods without the need for prior data integration.

## 3 Results

A typical workflow for TrAGEDy is as follows (Fig.1). Prior to trajectory alignment with TrAGEDy, a predefined list of marker genes of the various datasets being aligned is created; top variable genes or cluster markers can be selected, for example. The two datasets are then projected individually into a reduced dimension space using the preselected genes and TI performed on the two datasets independently (Fig.1A). There is no limitation on what TI method can be used, provided that pseudotime values are generated for each cell.

**Figure 1:**
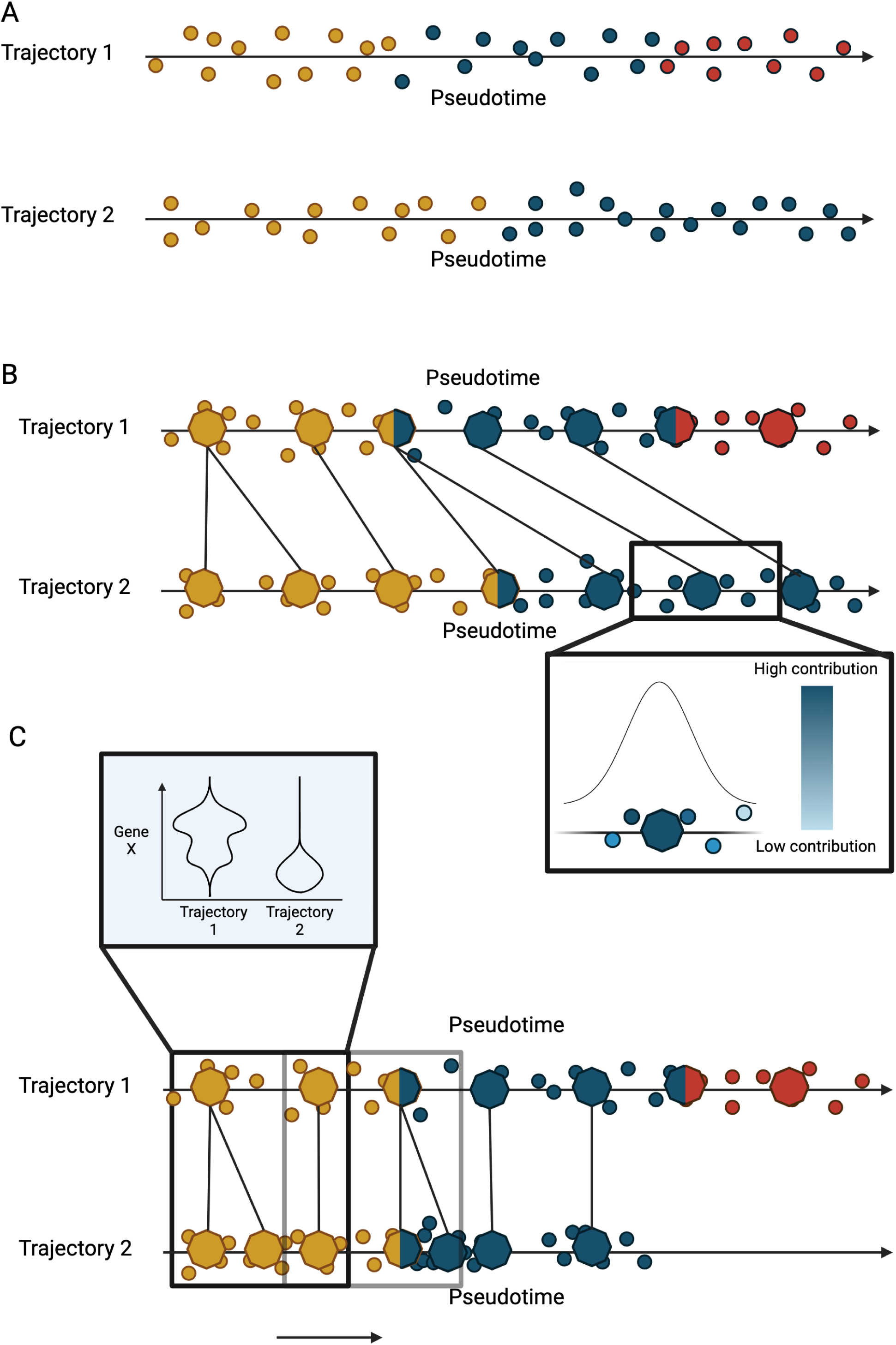
Graphical overview of the TrAGEDy process. Trajectory Inference is carried out on two datasets that share a common process but have a difference in condition (A). TrAGEDy samples gene expression across interpolated points of the trajectory, with cells with closer pseudotime values to the interpolated points contributing more to their gene expression (B). TrAGEDy then aligns the pseudotime of the interpolated points then the cells, finally performing a sliding window comparison between cells at similar points in aligned pseudotime, thereby extracting differentially expressed genes (C).

TrAGEDy then uses the cellAlign method of creating interpolated points to sample gene expression at different points in the trajectory, allowing inherently noisy scRNA-seq data to be smoothed across the trajectory. The closer a cell is to the interpolated point, the more it contributes to its gene expression. By changing a parameter that controls a gaussian pseudotime window, the user can alter the contribution level that more distal cells have to interpolated point gene expression (Fig.1B).

Dissimilarity between the gene expression of the interpolated points is next assessed. While interpolation can smooth out noise, possible batch effects between datasets mean calculating dissimilarity with methods such as Euclidean distance is problematic without scaling the data first. TrAGEDy uses Spearman correlation, as it is less sensitive to outliers than Pearson and, unlike Euclidean distance does not require scaling of gene expression values.

TrAGEDy next finds the optimal path through the dissimilarity matrix of the interpolated points, which constitutes the shared process between the two trajectories. DTW, with alterations, is used to find the optimal path. To overcome the constraint in DTW that the two processes must start and end at the same point, we first cut the dissimilarity matrix so the path starts and ends at points that are most transcriptionally similar to one another (supp. Fig.2). Another constraint of DTW is that all points must be matched to at least one other point; thus (post-DTW), we pruned any matches that have high transcriptional dissimilarity, enabling processes that may have diverged in the middle of their respective trajectories to be compared (supp. Fig.3). Finally, we used the modified alignment to adjust the pseudotime values of the interpolated points, generating new values as the basis to then adjust the pseudotime value of each cell (Fig.1C).

To extract DE genes, we performed a sliding window comparison, soft clustering cells at similar points in aligned pseudotime together and then testing for significance with a Mann-Whitney U test, extracting genes that are DE at different points in the shared process (Fig. 1C). Soft clustering with a sliding window allows comparisons over a continuous process. Users can define how much overlap there is between the windows of comparison.

### 3.1 Aligning simulated scRNA-seq datasets

We tested TrAGEDy’s ability to correctly identify the alignment of trajectories on datasets where the alignment points were known a priori. Using Dyngen (Cannoodt et al. 2021) we simulated scRNA-seq datasets with inherent trajectories encoded in the expression values, resulting in three different test cases: two positive controls and one negative control (Fig.2A).

**Figure 2:**
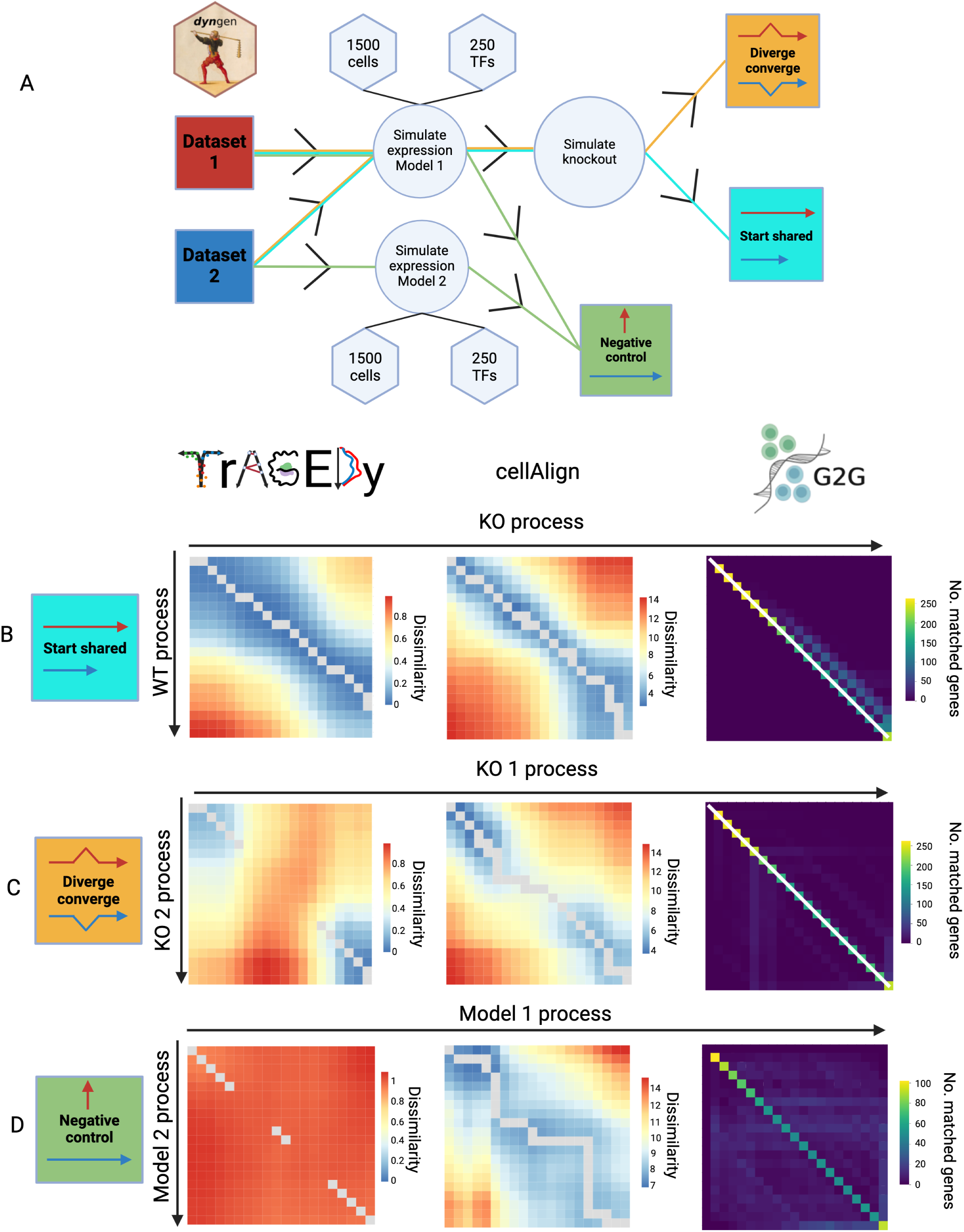
Alignment of simulated single cell RNA sequencing datasets with TrAGEDy and cell Align. Schematic showing the process of generating the three different topologies of alignment datasets, each with 1,500 cells and 250 transcription factors which drive the simulated process. For the diverge converge and start shared datasets, two datasets were generated using the same simulated expression model before knockouts were simulated for these datasets. For the negative control dataset, the expression of the two datasets was simulated using two different models (A). Heatmaps showing the alignment of the start shared (B), diverge converge (C) and negative control (D) Dyngen simulated datasets as assessed by TrAGEDy, cellAlign and Genes2Genes (G2G). Each box on the heatmap represents the alignment score of an interpolated point of each of the two datasets. For TrAGEDy and cellAlign the score represents transcriptional dissimilarity (as assessed by Spearman’s correlation and Euclidean distance respectively), while G2G shows the number of genes whose expression patterns is matched between the two interpolated points. The grey line represents the path of optimal alignment for TrAGEDy and cellAlign, while in G2G the optimal alignment is represented by a white line. The alignment heatmap of G2G for the negative control dataset has been edited to remove the alignment line; reflecting its conclusion that the two datasets share no common alignment.

The first positive test case used two datasets, the first simulating a full developmental progression involving sequential expression of many genes (wild type; WT), and the second resulting from knockout (KO) of one of these genes (Fig.2A). Thus, the KO trajectory is truncated at the point at which the process requires the transcription of the knocked out gene. TrAGEDy correctly aligned these datasets: the WT and the KO datasets initially align with one another, but the KO finishes its full process before the WT. For comparison, we also applied cellAlign and Genes2Genes (G2G), another trajectory alignment method which mainly focuses on aligning single genes, but can also be used to align multiple genes at once (Sumanaweera et al. 2023). Like TrAGEDy, the cellAlign distance matrix revealed the initial alignment of the WT and KO, before the KO trajectory ends while the WT continues. However, cellAlign failed to position the correct end point of the KO trajectory relative to the WT due to running DTW on the whole dissimilarity matrix. G2G also returns an incorrect alignment, suggesting that the simulated WT and KO datasets have a 1:1 alignment, although the number of matched genes does decrease as the trajectories progress (Fig.2B).

To assess how the methods handle a scenario where the trajectories have shared start and end points, but deviate in the middle, we simulated two datasets with Dyngen that have a diverge-converge backbone. By knocking out one of the divergent branches in one dataset (KO 1) and the other divergent branch in the other dataset (KO 2), we simulated linear trajectories that do not share a common process in their middle sections (Fig.2A). Applying TrAGEDy to this scenario accurately captured the initial and terminal alignment of the two trajectories, while leaving the middle sections unaligned. cellAlign failed to align KO 1 and KO 2 correctly, as cellAlign does not include functionality to prune matches. G2G also returned an incorrect alignment, as it suggested that there was a 1:1 alignment between the two simulated KO datasets (Fig.2C) despite the fact that they two datasets are transcriptionally divergent in the middle of their processes.

For the negative control, two datasets with distinct gene regulatory networks and transcription kinetics models were generated (referred to as model 1 and model 2) (Fig.2A). As there is no shared process between the conditions, the dissimilarity scores are expected to be high, and no alignment should be found. The dissimilarity score calculated by TrAGEDy between model 1 and model 2 showed that most of the matches have a dissimilarity score close to 1, and thus do not share a common process. As TrAGEDy finds the optimal path by looking at the context of the whole dissimilarity matrix, even if the dissimilarity scores are high, it will still find a path, as seen in the negative control (Fig.2C). TrAGEDy will give a warning to the user if the median of the cut dissimilarity matrix scores is over 0.6 and, thus the user must decide how to interpret the resulting alignment in the context of the overall dissimilarity score of the path. CellAlign also creates a path through the dissimilarity matrix, which has a higher overall dissimilarity range than the other two datasets, with the Euclidean distance ranging from 7-14 (Fig.2D), while the other two datasets ranged from 4-14 (Fig.2B, C). If, instead, Pearson correlation is selected for cellAlign, it becomes clearer from the alignment heatmap that the two datasets share little transcriptional similarity (supp. Fig.5C). As G2G directly models insertion and deletions in its alignment algorithm it can determine that there is no common alignment between the two datasets and, thus, is the only method to show that there is no underlying alignment in its alignment path (Fig.2D).

These analyses show that TrAGEDy is capable of faithfully capturing the underlying alignments of simulated biological scRNA-seq datasets, where current methods fail.

### 3.2 WT vs ZC3H20 KO Trypanosoma brucei alignment

Simulated data represents a ‘best case’ scenario for benchmarking tools, and often lacks the complexity of real datasets. We thus decided to apply TrAGEDy to real scRNA-seq datasets where the underlying processes have been characterised. For our first application, we examined a distinct developmental process in the kinetoplastid parasite *Trypanosoma brucei* (*T. brucei*). In the bloodstream of its mammalian host, WT *T. brucei* parasites transition from replicative slender forms to non-replicative stumpy forms, through a quorum sensing mechanism (Reuner et al. 1997, Dean et al. 2009,Rojas et al. 2019, Matthews 2021). These stumpy forms are preadapted to survive within the tsetse fly, which acts a vector for parasite transmission between mammals (Rico et al. 2013). Mutant *T. brucei* where an RNA binding protein essential for slender to stumpy differentiation, *ZC3H20*, has been knocked out (*ZC3H20* KO) are unable to undergo this transition (Liu, Kamanyi Marucha, and Clayton 2020,Cayla et al. 2020) and fail to express stumpy associated genes. Briggs and colleagues (2021) used 10X Chromium to sequence two biological replicates of WT *T. brucei* parasites (WT01 and WT02) undergoing this transition *in vitro*, as well as *ZC3H20* KO failing to differentiate in the same conditions. The authors defined four main clusters, Long Slender (LS) A and B, and Short Stumpy (SS) A and B, to show the progression of the transition, with LS A being the start and SS B being the end. Thus, the final alignment is expected to show that the WT and the *ZC3H20* KO datasets share an initial common developmental process, but that the KO process is truncated or branched relative to the WT, as suggested by previous analysis using data integration and Slingshot TI.

PHATE (Moon et al. 2019) embeddings were generated independently for the three datasets: two WT replicates (WT01 and WT02) and the *ZC3H20* KO. The PHATE embeddings were used as the basis for Slingshot TI to calculate pseudotime values for each of the cells. Applying TrAGEDy to WT01 and WT02 returned an alignment that started at the same point, and ended just before the final endpoint, returning a nearly 1:1 alignment, as predicted of biological replicates (supp. Fig.6). We next used the aligned WT values to perform TrAGEDy alignment between the combined WT replicates and the *ZC3H20* KO dataset. The resulting alignment showed that the two process have an initial point of alignment but the *ZC3H20* KO process was truncated relative to the WT (Fig.3A). Using the DTW calculated path, we then adjusted the pseudotime values of the trajectories to get a common pseudotime axis (Fig.3B).

**Figure 3:**
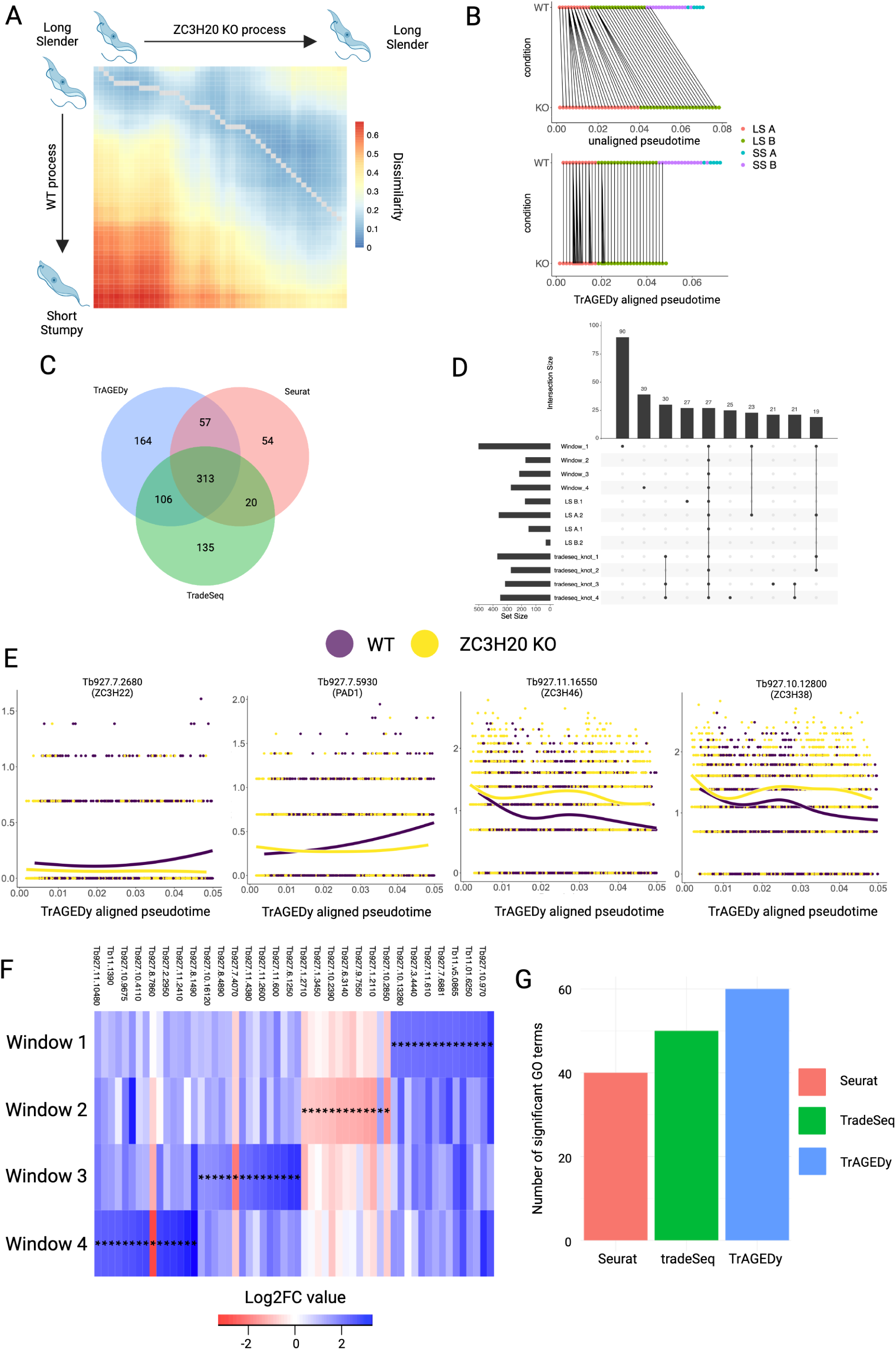
Analysis of WT and*ZC3H20* KO*Trypanosoma brucei* development with TrAGEDy. TrAGEDy alignment of the WT and *ZC3H20* KO trajectories of *Trypanosoma brucei* development with transcriptional dissimilarity calculated with Spearman correlation (A). TrAGEDy alignment of the interpolated points of the WT and *ZC3H20* KO trajectories, showing connections between the two processes. The interpolated points are displayed across pseudotime before and after the pseudotime values have been modified by TrAGEDy (B). Venn diagram showing the intersection of the differentially expressed (DE) genes captured by TrAGEDy, TradeSeq and Seurat (C). UpSet plot showing the top 10 intersections of differentially expressed genes captured by TrAGEDy across 4 windows of comparison and Seurat over 4 clusters and TradeSeq over 4 knot comparisons (D). TradeSeq plotSmoother plots showing the expression changes that occur across the WT (yellow) and *ZC3H20* KO (purple) trajectories within the cells of aligned pseudotime, as calculated by TrAGEDy, which share a common process. The 4 genes plotted were only identified as being significantly DE by TrAGEDy (E). Heatmap showing DE genes with the highest log 2 fold-change (Log_2_FC) captured (but not uniquely captured) between the WT and *ZC3H20* KO trajectories at 6 different windows of comparison. Each index is coloured by the Log_2_FC (red = higher expression in KO, blue = higher expression in WT) and a * symbol indicates the difference is significant (bonferroni adjusted p < 0.05, Mann-Whitney U test, abs(Log_2_FC) > 0.5, minimum percentage expressed > 0.1). A full list of heatmap genes can be found in supplementary table 7 (F). Barplot showing the number of significant Gene Ontology (GO) terms (Benjamini-Hochberg adjusted p-value < 0.01) returned when GO enrichment analysis was carried out using TriTrypDB on all the DE genes returned by TrAGEDy, TradeSeq and Seurat.

By aligning trajectories rather than integrating the datasets, we expected to preserve more variability between them, which may be reflected in the DE genes captured. We therefore compared TrAGEDy against other methods that can extract DE genes between conditions. We compared the differential gene expression tests by TrAGEDy with established methods of extracting DE genes: we compared TrAGEDy to a standard Seurat V5 pipeline (Y. Hao, Stuart, et al. 2024) of comparing DE genes between conditions, and TradeSeq’s method of comparing DE genes across trajectories from different conditions. Differentially expressed genes were defined as genes whose absolute log 2 fold change (Log_2_FC) was more than 0.75, Bonferroni adjusted p-value was less than 0.05, and were expressed in at least 10% of the cells in one of the comparison groups. To ensure comparability between the methods, the Log_2_FC of a gene between conditions was calculated using the same formula across all the different methods, namely the formula used in Seurat V5.

TrAGEDy detected more significant DE genes than both Seurat and TradeSeq (Fig.3C, D): TrAGEDy uniquely captured 164 genes as being DE, while TradeSeq and Seurat only returned 135 and 54, respectively (Fig. 3C, supp.table 1, 2, 3). Furthermore, the genes captured only by TrAGEDy that were more highly expressed in the WT were consistent with the slender to stumpy transition. These included the Arginine Kinase *AK3* (Tb927.9.6210) (Ooi et al. 2015), RNA binding proteins *ZC3H22* and *RBP38* (Tb927.7.2680, Tb927.11.5850) (Erben et al. 2021, Sbicego et al. 2003), Protein Associated with Differentiation 1 (*PAD1* - Tb927.7.5930) (Dean et al. 2009) and delta-1-pyrroline-5-carboxylate dehydrogenase (Tb927.10.3210) (Fig.3F, supp. table 1). For *ZC3H20* KO associated genes, two zinc finger genes, *ZC3H38*, *ZC3H46*, (Tb927.10.12800, Tb927.11.16550) (Lueong et al. 2016) were found to be differentially expressed towards the end of the shared process. Plotting the smoothed expression of some of these DE genes unique to TrAGDEy showed that the expression patterns matches the TrAGEDy output (Fig.3E). In terms of runtime required to return DE results, both TrAGEDy and Seurat returned results in under a minute, while TradeSeq took around 2 hours (supp. table 4).

Returning more significant DE genes and attaching biological meaning to individual genes are not robust metrics for assessing how well a DE gene detection method is performing, as they are prone to the influence of bias. As an example, randomly assigning 75% of the genes in the dataset to be DE would end up with more DE genes returned than most DE gene detection methods. Furthermore, there is no direct ground truth for whether the genes captured are DE or not. To address these issues, we used the number of significant GO terms returned from each methods DE gene list as a proxy for how biologically meaningful the output of the DE gene detection methods were likely to be. This approach was previously utilised in a previous analysis (Song and Li 2021), to test the usefulness of different DE gene detection methods (Song and Li 2021). We ran biological process GO term analysis on the three methods’ lists of DE genes, with GO term significance being set at Benjamini Hochberg procedure adjusted p-value < 0.01. Looking at the GO terms generated from the DE gene lists of the three methods showed that TrAGEDy returned 60 significant GO terms compared to 50 from TradeSeq and 40 from Seurat (Fig.3G) (Supp. table 5), with all but three of the GO terms returned by Seurat also returned by TrAGEDy and only 15 GO terms being unique to TradeSeq (supp. Fig.7B). When GO terms were generated on the list of uniquely captured genes for each method, TrAGEDy was the only method to return any significant GO terms (supp. Fig.7A, supp. table 6).

The above results show that TrAGEDy reveals more biologically relevant information on the slender to stumpy transitions under WT and *ZC3H20* KO conditions than established protocols.

### 3.3 WT vs *Bcl11b* KO T cell development analysis

Through applying TrAGEDy to the *T. brucei* datasets, we replicated an analysis that has already been carried out. With our next application, we used TrAGEDy in an analysis scenario which has not been attempted. For this we applied TrAGEDy to a scRNA-seq dataset of *in vitro* T cell development in WT and *Bcl11b* KO mice over two timepoints (Zhou et al. 2022). The data contain cells undergoing the early stages of T cell development in the thymus, from thymus-seeding precursors (TSP) to around the Double negative (DN) 4/Double positive (DP) stage. The authors reported that when *Bcl11b* expression is knocked out, the cells deviate from the T cell commitment pathway and develop distinct transcriptomic signatures around the DN2 stage of T cell development. However, some *Bcl11b* KO cells still express markers of T cell pathway commitment, such as the gene for the pre-T cell Antigen receptor α (*Ptcra*) (Hwang et al. 2020) and *Rag1* (supp. Fig.8), one of the central genes responsible for T cell receptor gene recombination, suggesting there may be some KO cells that have reached the post DN2 stages where the cells express a pre-TCR receptor and undergo β-selection (Mombaerts et al. 1992). In this context, we used TrAGEDy to compare the WT and *Bcl11b* KO trajectories of T cell development and attempted to identify differences in transcriptomes that occur between the WT and *Bcl11b* KO as they continue down the normal path of T cell development.

The cells were sequenced over two runs, with each run containing a variety of biological and technical replicates, as well as cells from different sampling days. For simplicity, the WT cells sequenced in run 1 will be referred to as WT1, the *Bcl11b* KO cells sequenced in run 1 will be referred to as *Bcl11b* KO1, and those sequenced in run 2 as WT2 and *Bcl11b* KO2, resulting in four individual datasets. When projected into the PHATE space, some of the WT1 cells were separated from the main body of the trajectory, which had downstream effects on the alignment, and so these cells were removed because TrAGEDy cannot function when there are gaps in the pseudotime axis (supp. Fig.9). TrAGEDy was then used to first align the two datasets for each condition together, resulting in two separate alignments: a WT alignment of WT1 and WT2, and a *Bcl11b* KO alignment of *Bcl11b* KO1 and *Bcl11b* KO2 (supp. Fig.10). The WT1 dataset only contains cells from day 10 of sampling, while WT2 contains cells from day 10 and 13. TrAGEDy captures this difference, showing a strong initial alignment of the two WT datasets with the WT1 dataset finishing before the WT2 (supp. Fig.10A). TrAGEDy was then performed on the aligned WT and *Bcl11b* KO trajectories. The resulting final alignment shows the two conditions sharing an initial common process, but the *Bcl11b* KO developmental progression finishes around the point when cells transition to express both TCR α and β chain genes (Fig.4A, B).

**Figure 4:**
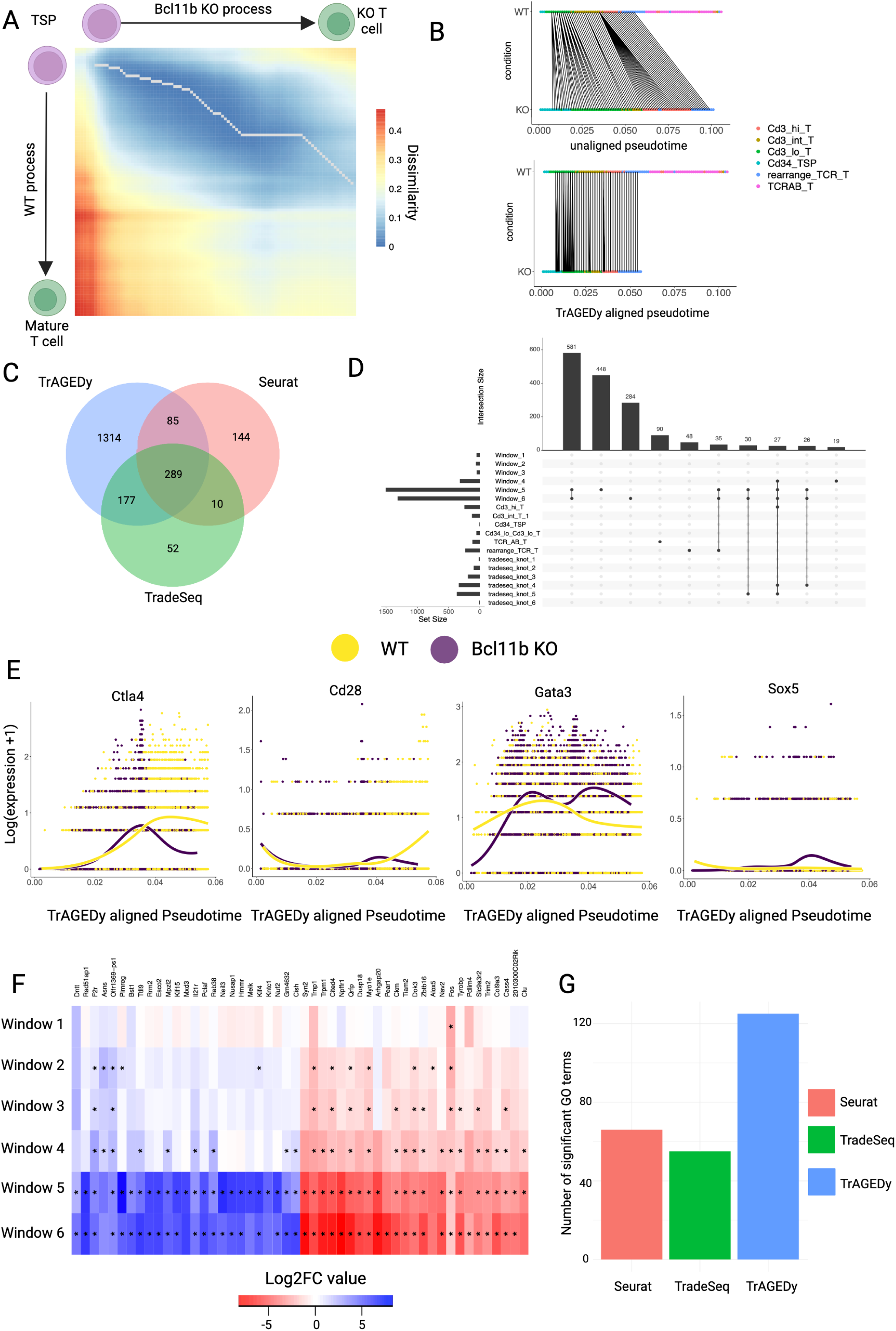
Analysis of WT and *Bcl11b* KO trajectories of T cell development with TrAGEDy. TrAGEDy alignment of the WT and *Bcl11b* KO trajectories of T cell development with transcriptional dissimilarity calculated with Spearman correlation (A). TrAGEDy alignment of the interpolated points of the WT and *Bcl11b* KO trajectories, showing connections between the two processes. The interpolated points are displayed across pseudotime before and after the pseudotime values have been modified by TrAGEDy (B). Venn diagram showing the intersection of the differentially expressed (DE) genes captured by TrAGEDy, TradeSeq and Seurat (C). UpSet plot showing the top 10 intersections of differentially expressed genes captured by TrAGEDy across 6 windows of comparison and Seurat over 6 clusters and TradeSeq over 6 knot comparisons (D). TradeSeq plotSmoother plots showing the expression changes that occur across the WT (yellow) and *Bcl11b* KO (purple) trajectories within the cells of aligned pseudotime, as calculated by TrAGEDy, which share a common process. The 4 genes plotted were only identified as being significantly DE by TrAGEDy (E). Heatmap showing DE genes captured (but not uniquely captured) by TrAGEDy with the highest log 2 fold-change (Log_2_FC) between the WT and *Bcl11b* KO trajectories at 6 different windows of comparison. Each index is coloured by the Log_2_FC (red = higher expression in KO, blue = higher expression in WT) and a * symbol indicates the difference is significant (bonferroni adjusted p < 0.05, Mann-Whitney U test, abs(Log_2_FC) > 0.5, minimum percentage expressed > 0.1). Full list of genes can be found in supplementary table 13(F). Barplot showing the number of significant Gene Ontology (GO) terms (Benjamini-Hochberg adjusted p-value < 0.01) returned when GO enrichment analysis was carried out using clusterprofiler on all the DE genes returned by TrAGEDy, TradeSeq and Seurat.

Overall, TrAGEDy returned 1865 DE genes across 6 windows of comparison, while TradeSeq and Seurat captured 528 DE genes across 6 knots and 6 cluster comparisons, respectively (Fig.4C, Supp. Table 8, 9 & 10). All the methods found more DE genes towards the end of the biological process compared with the beginning (Fig.4D). For genes only captured by one of the methods, TrAGEDy uniquely captured 1314 DE genes, while Seurat returned 144 and TradeSeq 52 (Fig. 4C, Supp. Table 8, 9 & 10). Genes with higher expression in the WT that only TrAGEDy identified as being DE include those associated with T cell signaling (*Cd28*, *Ctla4*) (Sansom 2000), cell cycle associated genes (*Top2a*, *Dut*, *Cenpa*) (Giotti et al. 2019), and those with functions for protecting against reactive oxygen species (ROS) (*Prdx4*, *Prdx1*) (Rhee et al. 2012). The genes only TrAGEDy identified as being significantly higher expressed in *Bcl11b* KO cells include pro-apoptotic factors (*Ikbip*, *G0s2* and *Stk4*) (Hofer-Warbinek et al. 2004, Welch et al. 2009, Cinar et al. 2007) and transcription factors (*Gata3* and *Batf*) (Wan 2014, Betz et al. 2010) (supp. Table 8). Plotting some of the DE genes that only TrAGEDy captured with TradeSeq smoothed expression show they are concurrent with the patterns of expression TrAGEDy suggests, with *Cd28* and *Ctla4* being more highly expressed in WT T cells, while *Gata3* and *Sox5* are more highly expressed in *Bcl11b* KO cells (Fig.4E). The times taken to return DE results was longer than the *T. brucei* analysis for all methods, however Seurat still managed to return results within seconds and TrAGEDy only took just over two minutes. In contrast, TradeSeq took over 20 hours to return a result (supp. table 4).

TrAGEDy found the most significant GO terms compared to Seurat and TradeSeq (Fig.4G, supp. Table 11). Furthermore, when only the uniquely captured DE genes were utilised, TrAGEDy was the only method that returned any significant GO terms (supp. Fig.11A, supp. Table 12). Of the significant GO terms returned by the methods for their entire DE gene list, only 14 were shared between all three methods (supp. Fig.11B), with many of them being GO terms related to regulation of lymphocyte development and activation (supp. Table 11). Of the GO terms that only TrAGEDy captured, most of them were related to regulation and activation of cell cycle-associated processes (supp. Table 11 & 12).

## 4 Discussion

In this paper we present TrAGEDy, a tool for aligning and comparing single cell trajectories between conditions. TrAGEDy allows the alignment of processes that start and end at different points by identifying the point at which dissimilarity increases from the minimum dissimilarity point. TrAGEDy assesses alignments to remove matches with high transcriptional dissimilarity in portions of the trajectories, allowing the algorithm to additionally capture alignments where the processes diverge and/or converge. TrAGEDy then identifies differences in gene expression over pseudotime between the conditions by taking a sliding window soft clustering approach, allowing it to identify transient changes in gene expression over the shared process.

Across multiple simulated datasets, TrAGEDy can find the underlying alignment between two processes, where cellAlign and G2G fails, due to its ability to prune matches and identify optimal start and end points in the dissimilarity matrix (Fig.2A). Across biological replicates and different conditions of real datasets, TrAGEDy overcomes batch effects to deliver accurate alignments and identify biologically relevant genes where TradeSeq and Seurat fail.

As of 2019, there were more than 70 TI methods, all of which have varying performance, functionality and constraints (Saelens et al. 2019). Additionally, the chosen method of dimensionality reduction (PCA, UMAP, PHATE, etc) used to generate the embeddings that the trajectory is calculated from also affects the pseudotime results. We chose PHATE for the reduced dimension embeddings as its purpose is to identify transitions in scRNA-seq data. User discretion is advised when choosing the TI and dimensionality reduction method as different methods will give different cell pseudotime values. The method of dimensionality reduction is especially important as many popular methods (like UMAP) have been found to distort cell relationships in the embedding space (Chari and Pachter 2023). Given the variability that can occur before TrAGEDy is applied, a core assumption of TrAGEDy is that the pseudotime must be an accurate reflection of the underlying biological process.

In almost all of the simulated datasets discussed here, TrAGEDy was able to correctly align trajectories, and with greater accuracy than cellAlign and G2G. For cellAlign, the constraints of DTW hamper it from accurately capturing the underlying alignments, while for G2G it may be due to the fact that the final alignment is based off of an aggregate of single gene alignments, rather than directly looking at the global alignment of genes across the cells. Additionally, TrAGEDy alignment of real data sets resulted in expected outcomes, including the complete alignment of biological replicates and partial alignments between conditions.

To compare DE results of TrAGEDy we also performed differential gene expression tests using two widely used tools: Seurat and TradeSeq. Although Seurat does not take pseudotime into account when it performs differential expression tests it is possible to compare cells across conditions, something which is not possible in many tools designed to specifically analyse trajectories, including pseudotimeDE and Monocle (Song and Li 2021, Trapnell et al. 2014). Low detection of DE genes by Seurat relative to TrAGEDy could be explained by the fact Seurat does not take pseudotime into account when carrying out the differential expression tests.

TrAGEDy identifies many known key processes which are required for *T. brucei* survival in the tsetse fly. While bloodstream forms do not wholly rely on glycolysis for its energy needs, most of its energy is produced through this pathway (Durieux et al. 1991, Taleva et al. 2023). In contrast to mammals, the tsetse fly is a low glucose environment (Y. Qiu et al. 2018), and the stumpy form is preadapted to suit these low glucose conditions (Grinsven et al. 2009). In the tsetse fly, the stumpy derived procyclic forms can breakdown amino acids for energy, in particular proline (Evans and Brown 1972, Weelden et al. 2003, Lamour et al. 2005). The parasite converts proline into glutamate through the action of proline dehydrogenase and 1-pyrroline-5-carboxylate dehydrogenase (*TbP5CDH*) with glutamate being further converted, through the action of glutamate-dehygrogenase, into 2-oxoglutarate, which can be used as an energy source in procyclic forms (Weelden et al. 2003, Mantilla et al. 2017). Only TrAGEDy captures *TbP5CDH* as being significantly upregulated in the WT at the end of the shared process, with TrAGEDy and Seurat both capturing glutamate dehydrogenase as being significantly upregulated in the WT towards the end of the shared process. None of the methods captured proline dehydrogenase as being DE at any point in the shared process. TrAGEDy thus paints a more complete picture of the proline catabolism pathway, which is active in the preadapted stumpy forms, than Seurat or TradeSeq. One of the key unique findings of TrAGEDy was identifying *PAD1* as being significantly upregulated in the WT towards the end of the shared process. *PAD1* is a key marker of stumpy forms, with its absence inhibiting induced differentiation of the parasite from stumpy to the first tsetse fly stage form, the procyclics (Dean et al. 2009).

When analysing the T cell development dataset, TrAGEDy captured more significant DE genes than Seurat and TradeSeq. While getting a higher number of DE genes is an important benchmark, it is also important to assess whether these genes fit into the context of the existing literature surrounding the datasets. TrAGEDy uniquely captured many cell-cycle associated genes as significantly upregulated in the WT compared with *Bcl11b* KO. After β-selection, before they begin to rearrange their TCR α chain, T cells undergo a proliferative burst and then become quiescent (Kreslavsky et al. 2012, Hwang et al. 2020). In accordance with this, cell cycle genes are mostly DE and upregulated in the WT in the final two windows (windows 5 and 6). Furthermore, the GO terms that only TrAGEDy captured were mainly associated with cell cycle function. This indicates that TrAGEDy has managed to capture the transient changes in proliferative state as WT T cells pass β-selection and prepare to rearrange their TCR α chain. Furthermore, TrAGEDy uniquely detected an increase in expression of *Cd28* and *Ctla4* in the WT compared to the *Bcl11b* KO. This finding further solidifies the conclusion that the WT T cells are able to pass β-selection while the *Bcl11b* KO are unable to, as *Ctla4* expression increases after T cell activation and an increase in *Cd28* expression has been seen following successful β-selection in maturing T cells (Linsley et al. 1992,Teague et al. 2010).

In terms of runtime, Seurat performed the best thanks to its optimisation through the Presto algorithm (Korsunsky et al. 2019) but TrAGEDy was not far behind in terms of runtime and performed better than Seurat in terms of the results returned. Tradeseq by performed the worst in terms of runtime, in some cases taking close to a day to return a result, due to the time needed to fit a general additive model to each gene in the dataset. The benchmarking of runtime was done single threaded. TradeSeq can be run in parallel, however the availability of computational power with the capability to run parallel is not widespread. TrAGEDy thus serves as a better alternative requiring less time to run, less resources and provides more biological insightful results than TradeSeq.

Ultimately, TI represents an approximation of how a developmental trajectory might look and may not represent the actual ordering of the cells through the process. Adding extra temporal information from lineage tracing will help add more biologically validity to single cell data (Wagner and Klein 2020). TrAGEDy could thus take lineage trace-inferred time, rather than pseudotime, as its input to allow the analysis of such datasets, allowing it to remain relevant as more sophisticated single cell sequencing techniques are introduced. Furthermore, the advent of perturb-seq allows researchers to more easily generate and analyse the effect of genetic perturbations on processes (Schraivogel et al. 2020), providing multiple and diverse opportunities for TrAGEDy to be applied.

## 5 Materials and Methods

The following is an overview of the steps that TrAGEDy takes to analyse a dataset. A workflow diagram of the TrAGEDy process and its core steps can be seen in supplementary figure 1 and implementation of the method can be seen, in full, in the code contained on GitHub (https://github.com/No2Ross/TrAGEDy).

### 5.1 Create interpolated points

Performing alignment directly on every cell in a dataset would be computationally and time expensive and prone to noise. As such, a modification of the cellAlign method was used to smooth gene expression over the trajectory by creating a user-defined number of interpolated points that sample the gene expression patterns of surrounding cells at specific points in the process and perform alignment on them. Each interpolated point is given the same sized set window of pseudotime around it. Cells within that window contribute highly to the gene expression of the interpolated point, while ones further away contribute less. TrAGEDy, allows the user to specify a window size, but this value is then weighted by the density of cells around each interpolated point. Interpolated points with many cells around it will have a smaller window, while those with few surrounding cells will have a larger window (supp. Fig.1B).

### 5.2 Scoring dissimilarity

For every interpolated point in the two trajectories, we calculate the transcriptomic dissimilarity between it and all the interpolated points on the other trajectory and store it in a dissimilarity matrix. This can be done using any metric, e.g. Pearson correlation, Spearman correlation, Euclidean distance, given the right processing of the data (e.g. scaling prior of gene expression prior to Euclidean distance). For our experiments, Spearman correlation was used to assess dissimilarity. We change the correlation scores to be 1 minus the correlation score hence, all the dissimilarity scales start from 0 (i.e. least dissimilarity) (supp. Fig.1C).

### 5.3 Identifying the optimal path of the two trajectories

To find the optimal alignment between the interpolated points of the two trajectories, Dynamic Time Warping (DTW) was utilised. DTW requires that every point on the two processes is matched with at least one other point on the opposing process. It also requires that the first and last points of each process must be matched to one another. To tackle this constraint, TrAGEDy scans the first and last row and column of the dissimilarity matrix and identifies the indexes with the lowest score from the beginning and end. These become the initial start and ends points of the process. To make sure no valid matches are missed from the downstream analysis, TrAGEDy takes the following steps. First, TrAGEDy finds out how much the absolute dissimilarity changes across the row/column of the beginning and end changes as we move across them (supp. Fig.2A). The user then defines whether to find the median or mean of these points and this value is used as a cutting threshold. To give a value to each index to compare against the cutting threshold, TrAGEDy finds the absolute difference between the scores of each index in the chosen row/column and the current start/end point (supp. Fig.2B). If there is an index value which occurs before the current start point or after the current end point whose dissimilarity score is less than the threshold, it is considered as a possible start/end point (supp. Fig.2B). For each possible path, we remove the interpolated points that fall before the start point and after the end point before performing DTW (supp. Fig.2C). To see what paths give us the best dissimilarity scores, we bootstrap the dissimilarity scores of the matched interpolated points along the path (supp. Fig.2C). The user can set the number of iterations during bootstrapping but the number of samples taken during each iteration set as the length of the longest path. The mean bootstrapped dissimilarity scores for each path is then calculated.

To deal with the scenario where two processes might differ somewhere in the middle of their processes, TrAGEDy first finds two thresholds for the matched and unmatched indexes (supp. Fig.3A). The user chooses to use either the mean or median of the dissimilarity score of all the unmatched indexes and all the matched indexes. If a matched index’s dissimilarity score is closer to that of the unmatched threshold than the matched threshold then it is cut, otherwise it remains matched (supp. Fig.3B).

### 5.4 Align pseudotime of interpolated points and cells

The aim of aligning the pseudotime of the interpolated points is to give matched interpolated points similar pseudotime values (supp. Fig.1D). As we have two processes, we have two vectors:

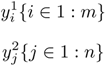

where m and n are the number of interpolated points on process 1 and 2 respectively, with 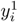 being the pseudotime of the *i* th interpolated point in process 1 and 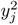 being the pseudotime of the *j* th interpolated point in process 2.

Interpolated points may only match one another or be a part of a multi-match, where one interpolated point on one of the trajectories is connected to 2 or more interpolated points on the other trajectory. For adjusting pseudotime of the former scenario (supp. Fig. 4A, Equation 1), given that 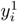 and 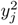 are matched and that 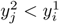:

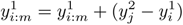

For a multi-match, first TrAGEDy aligns the pseudotime of the first section of the multimatch (supp. Fig.4B, equation 2), using the steps outlined for the individual match, then it does the same for the next match which isn’t in the multi-match (supp. Fig.4C, equation 3). The interpolated points that are multi-matched to one interpolated point on the other process, have their pseudotime values scaled between the aligned pseudotime value of the first match in the multi-match and the aligned pseudotime value of the match that occurs after the multi-match, adjusted by the difference between pseudotime of the next match and the pseudotime of the first match in the multi-match, normalized by the number of interpolated points being scaled (supp. Fig.4D, equation 4). This leaves us with an aligned pseudotime axis, where the matched interpolated points have similar pseudotime values (supp. Fig. 4E). We used the cellAlign method of mapping gene expression values from individual cells onto interpolated points to map pseudotime values from interpolated points onto the individual cells.

### 5.5 Differential expression analysis with TrAGEDy

TrAGEDy identifies differentially expressed (DE) genes across pseudotime, between the two conditions by taking a sliding window soft clustering approach (supp. Fig.1E).

First the cells were assigned to the closest interpolated point in terms of pseudotime via one round k-means clustering, with the interpolated points acting as the cluster centroid. The interpolated points that were matched together were grouped, with the cells that were assigned to these interpolated points acting as a cluster which the window slides over when making comparisons. The user defines how many windows of comparison will be made across the pseudotime, and how much overlap there will be, in terms of matched interpolated points, between each successive window.

The user chooses a method of assessing significance using either a t-test or a Mann-Whitney U test. Log_2_ fold change (Log_2_FC) of the genes between conditions is calculated using the same formula used by Seurat V5. To adjust the p-values to account for multiple testing, Bonferroni correction is used across all the genes in the dataset.

### 5.6 Simulating trajectories with Dyngen

All simulated datasets were simulated with Dyngen (Cannoodt et al. 2021). The first positive control experiment trajectories were simulated with a linear backbone with parameters drawn from the same transcription factor, feature and kinetic network distributions. One of the datasets was not modified beyond the standard pipeline, while the other had a full knockout in the B5 gene module, causing a stunted process in comparison to the other dataset. The second positive control experiment trajectories were simulated with a bifurcating-converging backbone with parameters drawn from the same transcription factor, feature and kinetic network distributions. One of the datasets had a full knockout in the C1 gene module while the other dataset had a full knockout in the D1 gene module. The negative control experiment trajectories were simulated with a linear backbone with parameters drawn from different transcription factor, feature and kinetic network distributions. The specific parameters and program calls used to simulate the individual datasets can be found on the GitHub link in the “Availability of data and materials” section.

### 5.7 Applying TrAGEDy, Genes2Genes and cellAlign to simulated trajectories

All simulated datasets were processed using the following. In Seurat, the simulated counts were normalized prior to scaling across all simulated genes. Principal Component Analysis (PCA) was performed on the scaled data and using the first 5 PCA dimensions clustering was performed at a resolution of 0.2 for all the datasets as well as UMAP. Slingshot (Street et al. 2018) was performed to get pseudotime values for each of the cells, using the UMAP space as the basis for the trajectory. To get the feature space, the top 100 DE genes in terms of Log_2_FC for each cluster in each dataset were collected for both datasets being compared. The prior analysis steps were kept the same for both cellAlign and TrAGEDy analysis, as well as the window size and the number of interpolated points. For cellAlign Euclidean distance and Pearson correlation was used to calculate dissimilarity, while for TrAGEDy Spearman correlation was used. For Genes2Genes, a binning number of 20 was used for all the comparisons and the same feature space was used as in the TrAGEDy and cellAlign analyses. More specific parameter values used for each dataset can be found on the TrAGEDY github, linked in “Availability of data and materials” section.

### 5.8 Pre-processing of WT vs *ZC3H20* KO *Trypanosoma brucei* dataset

The processed RDS (R dataset) file containing the final cluster labels was acquired from the authors. The ‘LS A.1’ and ‘LS A.2’ clusters were merged into the cluster ‘LS A’ and the ‘LS B.1’ and ‘LS B.2’ were merged into the cluster ‘LS B’.

### 5.9 TrAGEDy analysis of WT vs *ZC3H20* KO *Trypanosoma brucei* dataset

The integrated object was split into the individual datasets (WT01, WT02 and ZC3H20 KO). The top 200 DE genes (defined as absolute log_2_FC > 0.75, Bonferroni corrected p-value < 0.05 and the gene expressed in at least 25% of cells) across the clusters were found for each dataset and used as the basis for constructing the PHATE space and assessing similarity with TrAGEDy. The number of genes in the feature space amounted to 455 genes. PHATE embeddings were constructed from the normalised gene expression count matrix. Pseudotime was calculated using Slingshot for each of the three datasets, with the ‘LS A’ cluster used as the starting point. The WT01 and WT02 datasets were then aligned with TrAGEDy, with 50 interpolated points used to align the processes. This TrAGEDy aligned WT dataset was then aligned against the *ZC3H20* KO dataset. Differential expression was carried out over 4 windows, with a gene identified as being DE if the absolute Log_2_FC was more than 0.75, bonferroni corrected p-value < 0.05 and the gene expressed in at least 10% of cells in one of the conditions.

### 5.10 Seurat analysis of WT vs *ZC3H20* KO *Trypanosoma brucei* dataset

The ‘SS A’ and ‘SS B’ clusters were removed from the integrated dataset, as they are not present in the *ZC3H20* KO dataset, and the FindMarkers function was performed for all the four slender clusters (LS A.1, LS A.2, LS B.1 & LS B.2) between the WT and *ZC3H20* KO conditions. A gene identified as being DE if the absolute Log_2_FC was more than 0.75, bonferroni corrected p-value < 0.05 and the gene expressed in at least 10% of cells in one of the conditions.

### 5.11 TradeSeq analysis of WT vs *ZC3H20* KO *Trypanosoma brucei* dataset

The ‘SS A’ and ‘SS B’ clusters were removed from the integrated dataset. Using the authors PHATE space, Slingshot was applied to build trajectories. The optimal number of knots for fitting the general additive model to the dataset was assessed through the evaluateK function, with the optimal number determined to be 8. fitGAM was run on the dataset followed by the conditionTest function to identify DE genes between the WT and *Bcl11b* KO conditions. 4 knot comparisons were made, and a gene were determined to be DE if the absolute Log_2_FC was greater than 0.75, bonferroni p-value < 0.05 and the gene was expressed in at least 10% of the cells that fell within the current knot comparison, in one of the conditions.

For all cases, genes were analysed in TriTrypDB to get their gene short names and functions (Amos et al. 2022, Alvarez-Jarreta et al. 2024).

### 5.12 Preprocessing of WT vs *Bcl11b* KO *in vitro* T cell development dataset prior to TrAGEDy analysis

The RDS (R dataset) file containing the scRNA-seq information was acquired from the authors (Zhou et al. 2022).

The dataset was split into the WT and *Bcl11b* KO conditions and then further split into the different sequencing runs. For each of the four scRNA-seq objects derived from the previous, the data was analysed using the Seurat V4 pipeline. When the data was scaled, the effects of the cell cycle were regressed out by passing the cell cycle associated genes Seurat provides into the var.to.regress parameter. For each dataset, PCA was carried out on the scaled-regressed expression matrix, reducing the dimensions down to 50. An elbow plot was used to determine the number of Principal Components (PC) used to create a UMAP space embedding. Clustering was performed using the same number of PCs for UMAP embedding calculation. Clustree (Zappia and Oshlack 2018) was used to help choose a resolution for clustering that leads to stable clusters. Clusters were annotated as follows. Clusters that did not express Cd3 genes but expressed Cd34 were defined as ‘Cd34 TSP’, clusters which did not express Cd34 or Cd3 were characterized as ‘early T’, clusters which had low expression of Cd3 but no Cd34 expression were characterised as ‘Cd34 lo Cd3 lo T’, those with high Cd3 expression were ‘Cd3 T hi’ and those with middle levels of Cd3 expression were ‘Cd3 T mid’. Clusters which had high expression of Cd3 and expressed *Ptcra* and *Rag1* were defined as ‘rearrange TCR T’ and those that expressed the TCR signal transduction molecule *Zap70* and some cells that express the gene Trac, were defined as ‘TCR AB T’. Clusters that express Rora were defined as ‘Rora T’, clusters that expressed high levels of interferon associated genes were defined as ‘Interferon response’ and cells which clustered out with the main body of the UMAP were classed as ‘Outlier’. The ‘Interferon response’ and ‘Outlier’ clusters were removed before downstream analysis with TI, TrAGEDy, Seurat DE or TradeSeq.

### 5.13 TrAGEDy analysis of WT vs *Bcl11b* KO *in vitro* T cell development dataset

In order to build a gene feature space to construct PHATE embeddings and assess dissimilarity with TrAGEDy, the feature space for the dataset was made of the combined DE genes for each cluster whose absolute Log_2_FC > 0.5, Bonferroni corrected p-value < 0.05 and was expressed in at least 1%, across all four of the datasets. To reduce the effect of the cell cycle, genes supplied by Seurat as being important in the cell cycle and genes included in the Mouse Genome Informatics GO term ‘cell cycle’ were removed from the feature space. For all four of the individual datasets, 10 dimensional PHATE embeddings were generated from the normalised gene expression count matrix for the features selected previously. For the WT1 dataset, cells with a pseudotime higher than 0.1 (11 cells) were removed. TI was then carried out using Slingshot on the PHATE embeddings with the Cd34 TSP cluster chosen as the starting point. TrAGEDy was carried out on each of the conditions, using 100 interpolated points with a window size of the maximum pseudotime value, from either of the two datasets being analysed, divided by 90. TrAGEDy aligned pseudotime values were then created, resulting in a TrAGEDy aligned WT and a TrAGEDy aligned *Bcl11b* KO dataset.

The WT and *Bcl11b* KO TrAGEDy aligned datasets were then analysed using TrAGEDy. TrAGEDy was carried out with the same number of interpolated points and window size as the replicate alignments. TrAGEDy differential expression test was then carried out across 6 windows of comparison. A gene was returned as being differentially expressed if the absolute Log_2_FC was greater than 0.75, Bonferroni adjusted p-value < 0.05 and the gene was expressed in at least 10% of cells in the window of comparison, in one of the conditions. The final TrAGEDy aligned WT and *Bcl11b* KO datasets were then analysed independently by TradeSeq to create the smooth gene expression plots across the aligned pseudotime.

### 5.14 Seurat analysis of WT vs *Bcl11b* KO *in vitro* T cell development dataset

Using Seurat V5 integration (Satija et al. 2015, Butler et al. 2018, Stuart et al. 2019, Y. Hao, S. Hao, et al. 2021, Y. Hao, Stuart, et al. 2024), the four datasets were integrated using the ‘integrated.cca’ method. When the data was scaled, the effects of the cell cycle were regressed out by passing the cell cycle associated genes Seurat provides into the var.to.regress parameter. PCA was carried out on the scaled-regressed expression matrix, reducing the dimensions down to 50. An elbow plot was used to determine the number of Principal Components (PC) used to create a UMAP space embedding. Clustering was performed using the same number of PCs for UMAP embedding calculation. Clustree was used to help choose a resolution for clustering that leads to stable clusters and clusters were annotated as described previously. The FindMarkers function was used to identify DE genes between the conditions for each cluster. Genes were determined to be DE if the absolute Log_2_FC was greater than 0.75, bonferroni adjusted p-value < 0.05 and the gene was expressed in at least 10% of the cells in one of the conditions for the cluster being compared.

### 5.15 TradeSeq analysis of WT vs *Bcl11b* KO *in vitro* T cell development dataset

Using the Seurat integrated, cell cycle regressed dataset; PHATE was carried out, using the scaled and cell cycle regressed gene expression matrix. Slingshot was then applied to the dataset, using the Cd34 TSP cluster as the starting point of the trajectory. Cells on the trajectory path which ended with the TCR AB T cluster was kept and all other cells were removed from the dataset. The optimal number of knots for fitting the general additive model to the dataset was assessed through the evaluateK function, with the optimal number determined to be 7. fitGAM was run on the dataset followed by the conditionTest function to identify DE genes between the WT and *Bcl11b* KO conditions. 6 different knot comparisons were made, and a gene were determined to be DE if the absolute Log_2_FC was greater than 0.75, bonferroni p-value < 0.05 and the gene was expressed in at least 10% of the cells that fell within the current knot comparison, in one of the conditions.

### 5.16 GO term analysis of T cell DE genes

Biological process GO term analysis of the lists of DE genes (either unique DE genes or all of them) for the three methods was carried out using the enrichGO command of clusterProfiler. Background genes were the genes whose expression was more than 5% across the entire integrated T cell dataset. GO terms whose Benjamini-Hochberg corrected p-value was less than 0.01 were identified as significantly enriched GO terms.

### 5.17 GO term analysis of *Trypanosoma brucei* DE genes

Biological process GO term analysis of the lists of DE genes (either unique DE genes or all of them) for the three methods was carried out using TriTrypDB. GO terms whose Benjamini-Hochberg corrected p-value was less than 0.01 were identified as significantly enriched GO terms.

### 5.18 Runtime experiments

Packages were run single threaded on a virtual machine with 70 gigabytes of allotted RAM and 3 GHz clock speed. Runtime experiment code detailing what sections were included for each package when calculating runtime can be found on our GitHub.

### 5.19 Package versions details

All analysis and experiments were carried out on R were done with version 4.1.2 except for the Dyngen dataset simulation and the TradeSeq experiments which were carried out on R version 4.1.0. Seurat version 5.0.3, Dyngen version 1.0.5, phateR version 1.0.7, Slingshot version 2.2.1, Single Cell Experiment version 1.16.0, cellAlign version 0.1.0, zellkonverter 1.4.0 & TradeSeq version 1.6.0 were used.

For the Genes2Genes experiments, analysis was done on Python version 3.8.16 with the following package versions: genes2genes version 0.2.0, numpy version 1.24.3, pandas version 2.0.3, scanpy version 1.9.3, scipy version 1.10.1, matplotlib version 3.7.1, seaborn version 0.12.2 & scikit-learn version 1.2.2.

### 5.20 Availability of data and materials

The functions to run TrAGEDy are available on our GitHub (https://github.com/No2Ross/TrAGEDy) along with the code required to repeat our analysis.

## 6 Contributions

R.F.L. and T.D.O. conceived the project and R.F.L. implemented the TrAGEDy package. R.F.L., T.D.O., E.M.B. and R.M. designed experiments. R.F.L. performed the experiments and collected data. R.F.L., T.D.O., E.M.B. and R.M. analyzed the data. R.F.L. wrote the manuscript with editing from R.M., E.M.B., K.R.M. and T.D.O. All authors read, edited and approved the manuscript.

## 7 Ethics declarations

### 7.1 Ethics approval and consent to participate

Not applicable.

#### 7.1.1 Competing interests

The authors declared that they have no competing interests.

## 8 Funding

This research was funded by the Medical Research Council through their Precision Medicine DTP (MRC grant number: MR/N013166/1). This research was funded in whole, or in part, by the Wellcome Trust [Grant numbers 218648/Z/19/Z to EMB, 104111/Z/14/ZR to TDO/RM, 206815/Z/17/Z & 224501/Z/21/Z to RM, and 103740/Z14/Z to KRM] and the BBSRC Project grants (BB/R017166/1 & BB/W001101/1 to RM). TDO is further funded by the ExposUM Institute of the University of Montpellier.

## 9 Acknowledgements

The authors would like to thank Scott Arkison for maintaining the computational infrastructure.

## 11 Supplementary figures

**Supplementary Figure 1:**
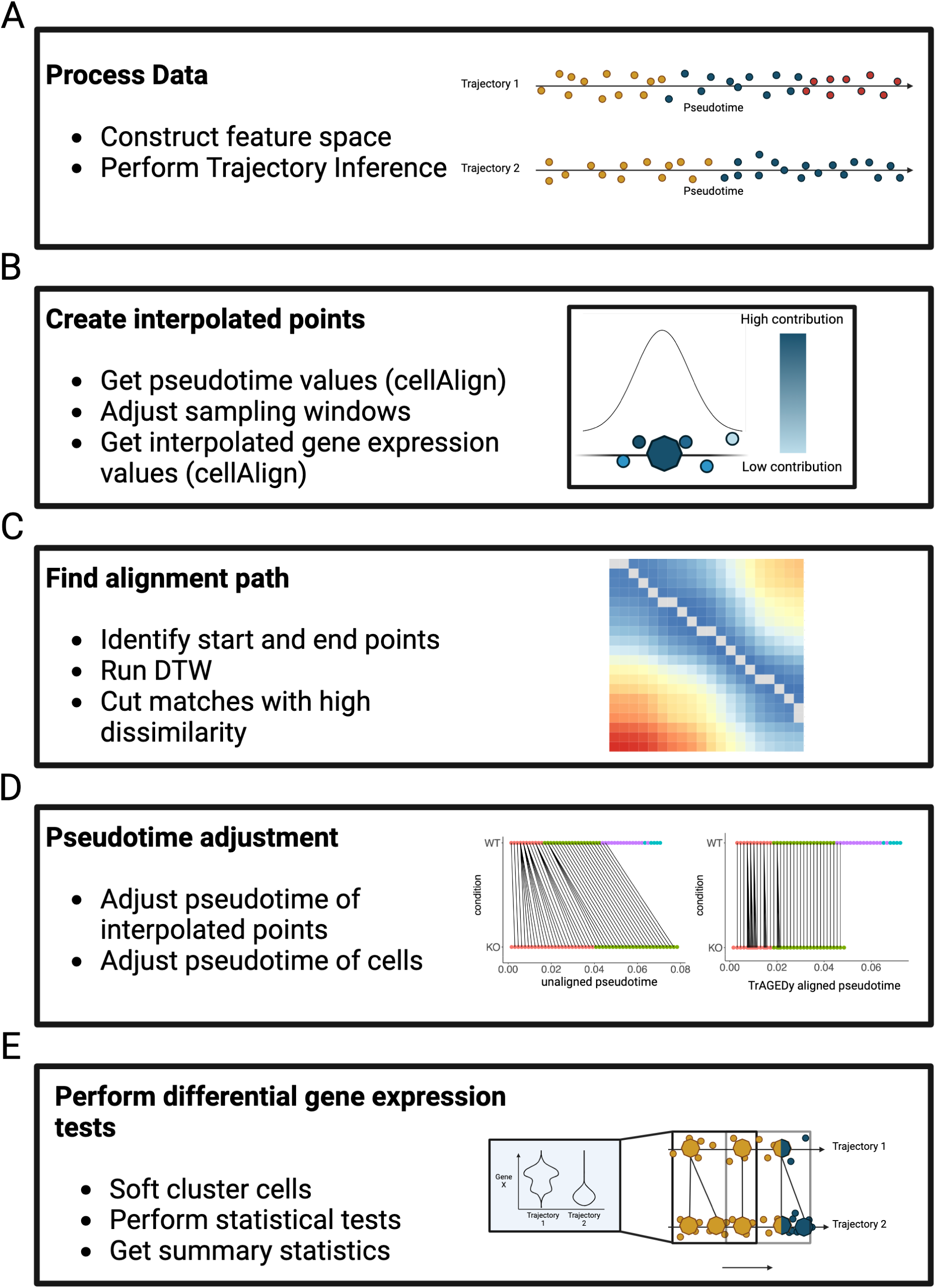
Workflow of TrAGEDy process. Workflow detailing the different parts of the TrAGEDy process. Sections which were derived from cellAlign are noted in the process boxes. TrAGEDy starts with the user performing Trajectory Inference (TI) on the datasets individually using a shared feature space (A). The pseudotime and gene expression of the cells is then used to create interpolated points using the method defined by cellAlign (B). From the interpolated points, TrAGEDy then works out the optimal alignment by first identifying the optimal start and end points before running DTW and pruning any suboptimal matches (C). The alignment path is then used to adjust the pseudotime of the interpolated points, and then subsequently the cells themselves (D). Differentially expressed genes are then identified across the shared process to see identify what gene expression changes occur before the datasets diverge from one another (E).

**Supplementary Figure 2:**
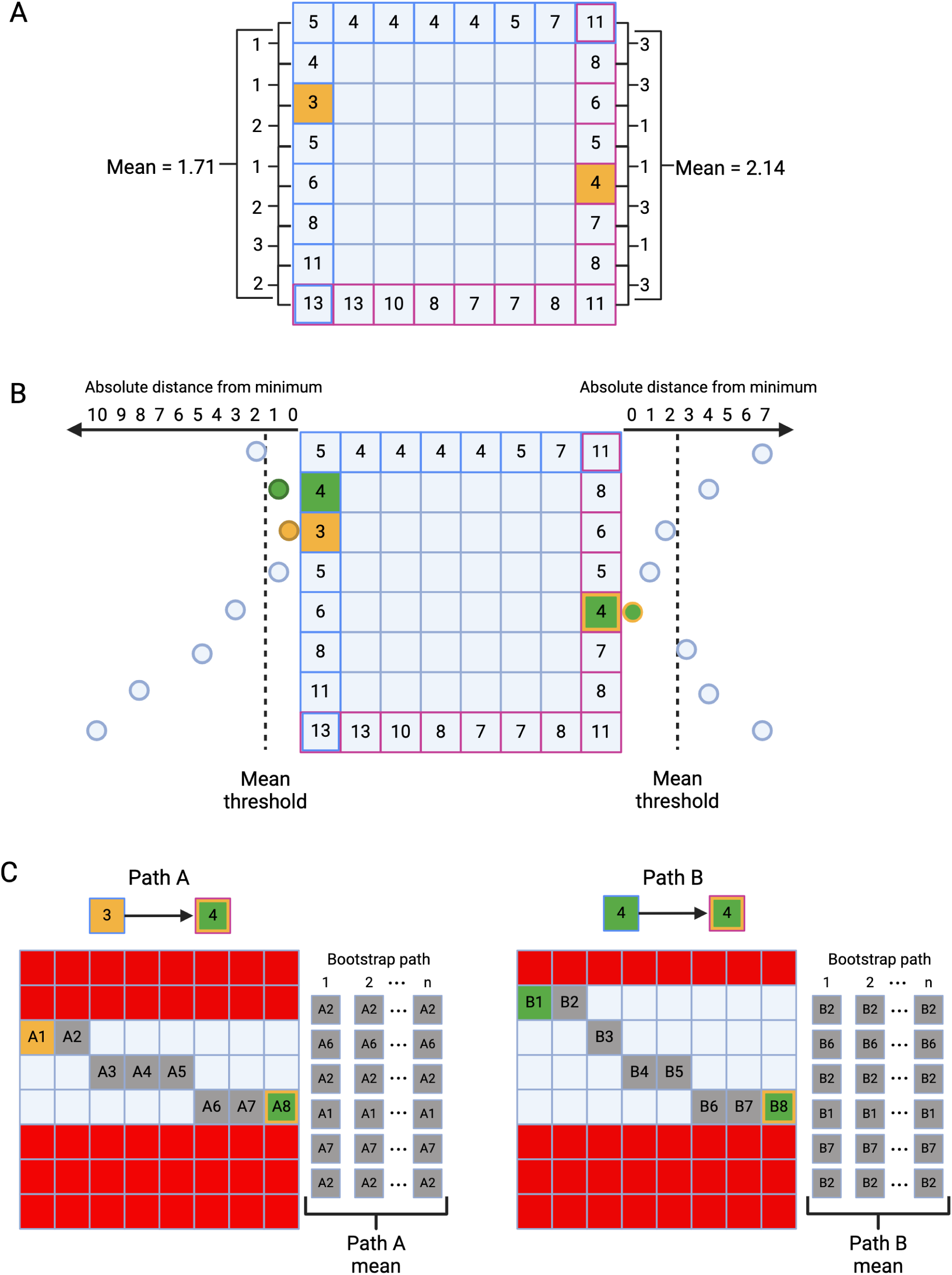
Graphical explanation of the TrAGEDy process of identifying the optimal start and end points. TrAGEDy identifies the optimal start and end points by first finding the slice of the score matrix from the start (outlined in blue) and end (outlined in purple) which contains the lowest dissimilarity score match (box coloured orange). TrAGEDy then finds the mean/medians of the score differences between adjacent matches on the start and end slice (A). The means/medians are then used as threshold for considering which matches are the optimal start and end points, with matches whose dissimilarity score is less than the threshold and before the current start match and after the current match (green box) are considered as possible start/end matches (B). For each of the possible paths through the data TrAGEDy performs the following steps. First, it removes any matches (shown in red) that fall outwith the select start and end points and then performs Dynamic Time Warping (DTW) on the cut score matrix. The matches included in the DTW path are then bootstrapped and the mean of the bootstrapped paths is calculated (C).

**Supplementary Figure 3:**
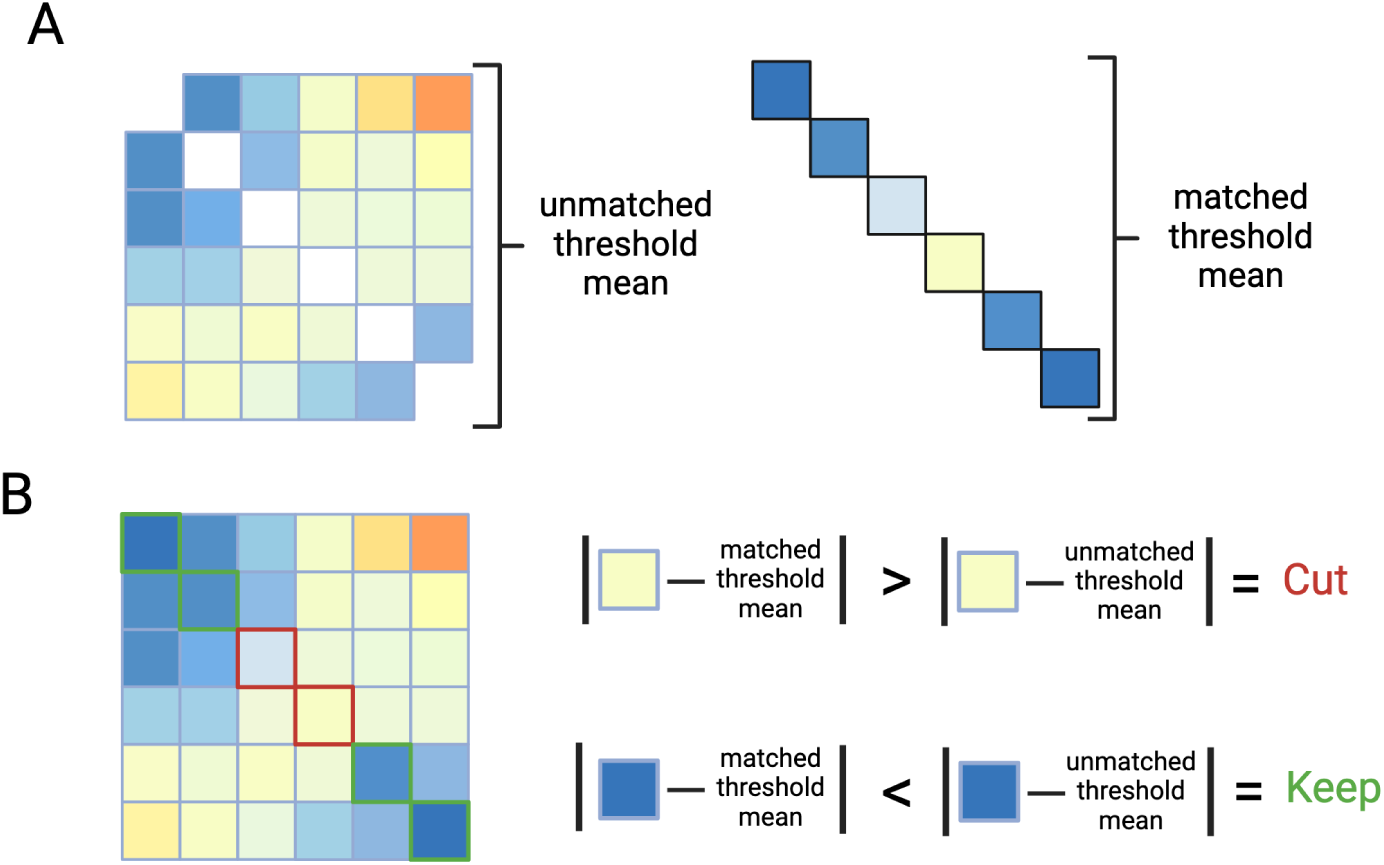
Graphical explanation of the TrAGEDy process of cutting highly dissimilar matches. To remove matches from the path which are not optimal TrAGEDy goes through the following process. An unmatched threshold and matched threshold are created by taking the mean/median of scores of the unmatched and matched points respectively (A). Simply, if the absolute difference between a matches score is closer to the unmatched threshold than the matched threshold it is cut and if the opposite is true, it is kept (B).

**Supplementary Figure 4:**
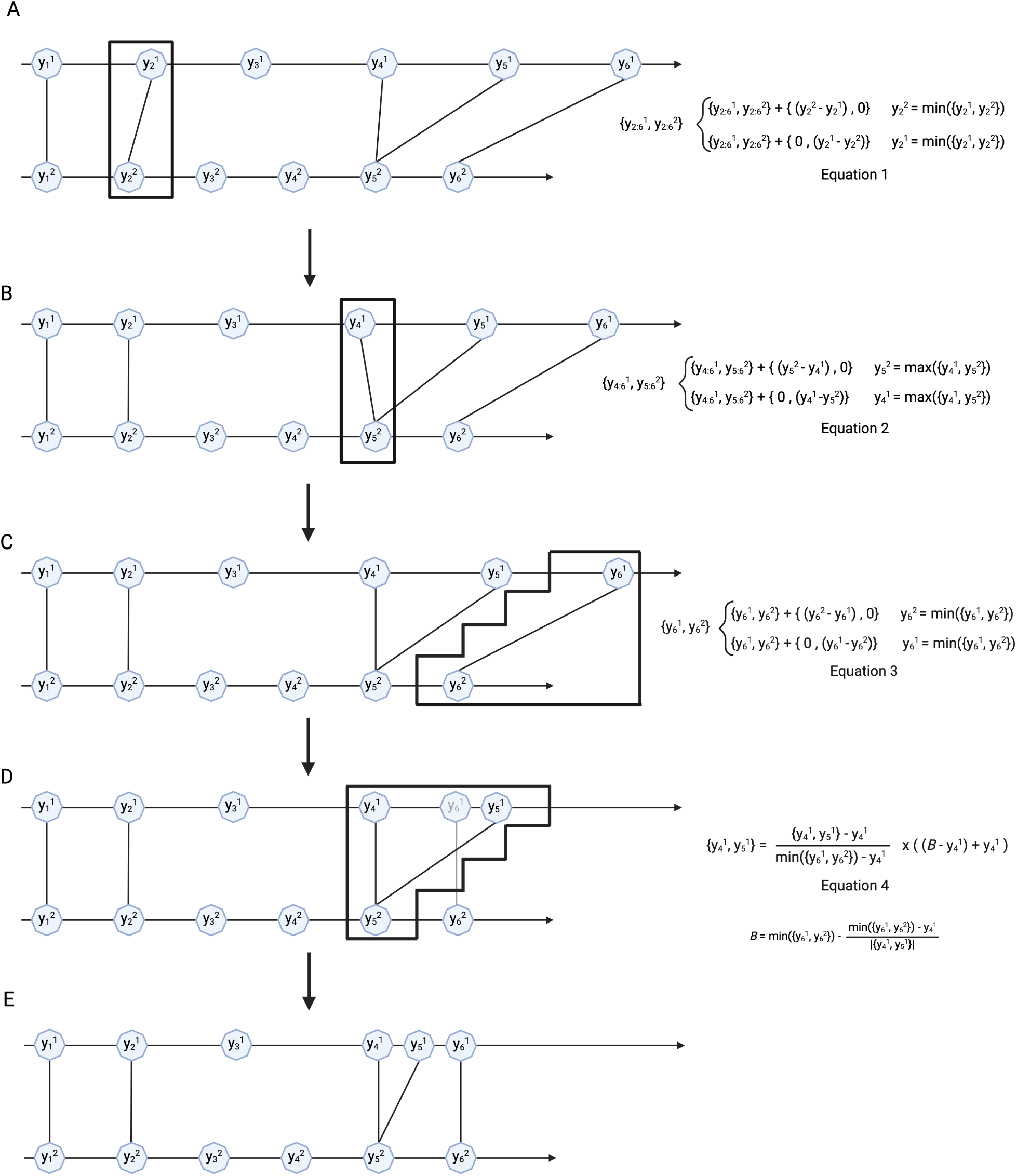
Graphical explanation and example of the alignment of interpolated point pseudotime values, from the DTW calculated path. There are two types of movement that can occur when aligning pseudotime values, an individual match (A, B & C) or a multi-match (D). For the individual match, the pseudotime value of the interpolated point with the highest value (and all the subsequent interpolated points) are brought down by the difference between the lower pseudotime value and the higher pseudotime value. For multi-match points, the match that occurs after it is adjusted then the first match in the multi-match. All the interpolated points that are matched to the same interpolated point are then scaled between pseudotime of the initial match in the multi-match and the pseudotime of the next match modified by the difference between the two, normalised by the number of interpolated points to be scaled. This then gives us aligned pseudotime values for all the interpolated points (E).

**Supplementary Figure 5:**
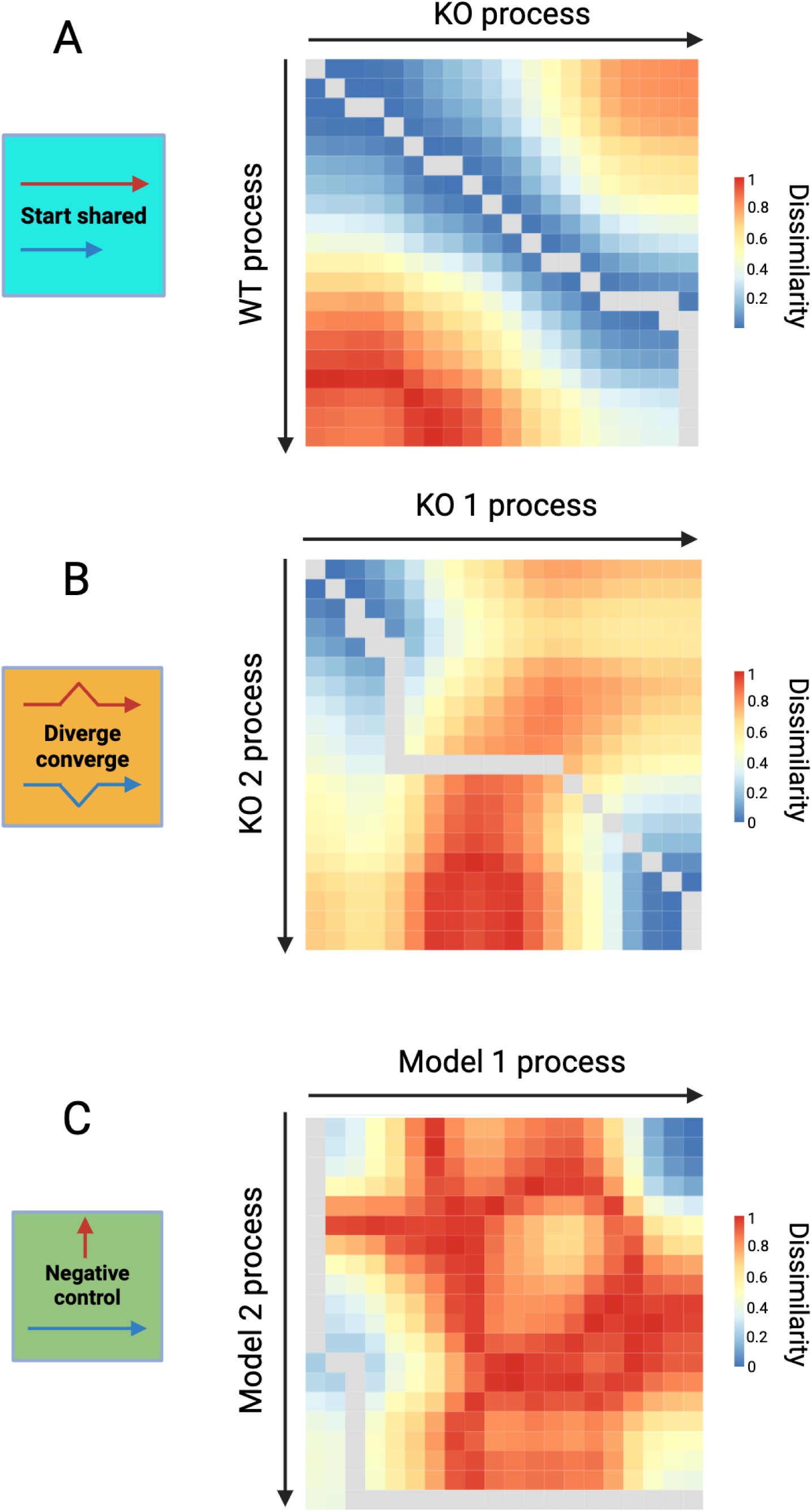
cellAlign alignments of simulated datasets using Pearson correlation dissimilarity score. Heatmaps showing the alignment of the two datasets across the three Dyngen simulated alignment topologies. Each box of the heatmap shows the transcriptional dissimilarity (as assessed by Pearson correlation) of interpolated points of the two datasets with blue meaning low dissimilarity and red meaning high dissimilarity. The grey line denotes the optimal alignment of the interpolated points.

**Supplementary Figure 6:**
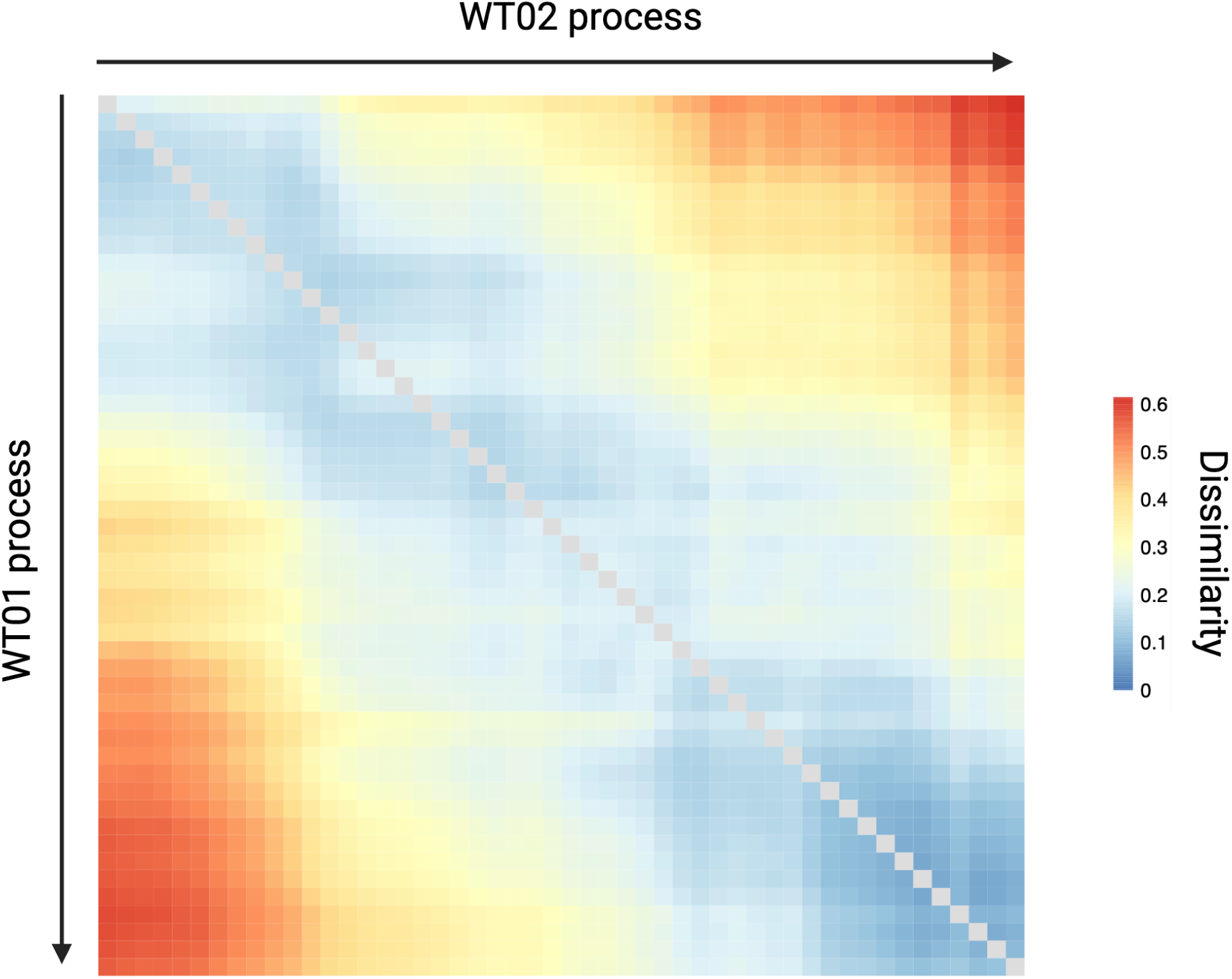
TrAGEDy alignment of biological replicates of WT slender to stumpy *Trypanosoma brucei* transition. TrAGEDy alignment of the WT01 and WT02 slender to stumpy *Trypanosoma brucei* development trajectories. Dissimilarity in gene expression of interpolated points was calculated using Spearman correlation with blue meaning low dissimilarity and red meaning high dissimilarity.

**Supplementary Figure 7:**
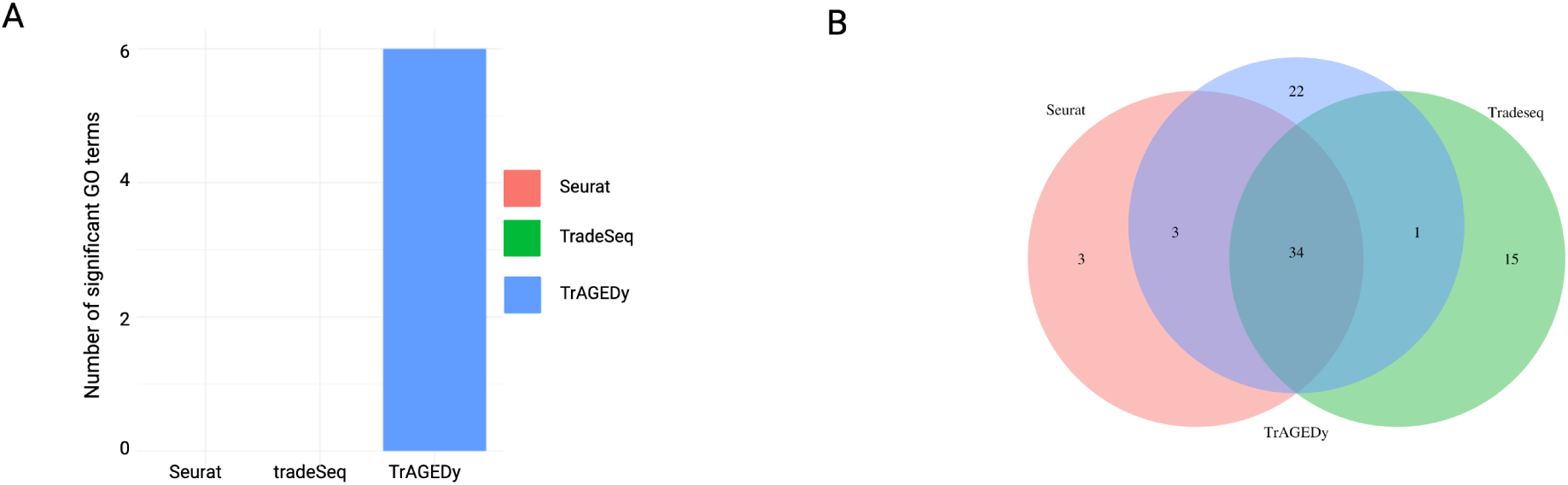
Further GO term analysis of WT vs *ZC3H20* KO *Trypanosoma brucei* bloodstream form development. Barplot showing the number of significant Gene Ontology (GO) terms (Benjamini-Hochberg adjusted p-value < 0.01) returned when GO enrichment analysis was carried out using clusterprofiler on the unique DE genes returned by TrAGEDy, TradeSeq and Seurat (A). Venn diagram showing the intersections of significant GO terms (Benjamini-Hochberg adjusted p-value < 0.01) found by the three methods using clusterprofiler on all the DE genes returned by TrAGEDy, TradeSeq and Seurat (B).

**Supplementary Figure 8:**
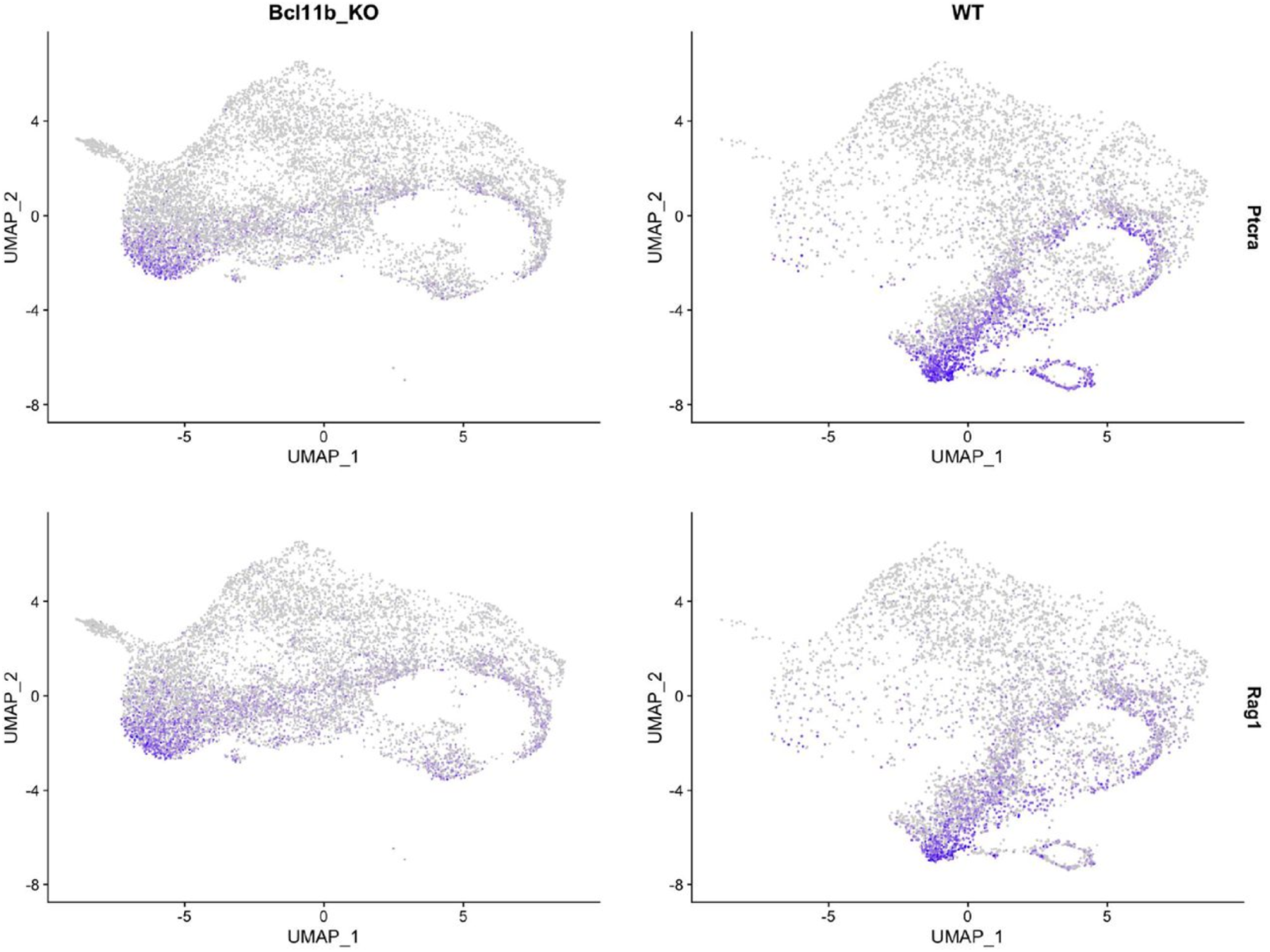
Expression of markers associated with DN3 stage of T cell development in *Bcl11b* KO and WT T cells. UMAP showing the normalised gene expression levels of *Ptcra* (pre-T cell receptor α chain) and *Rag1* (recombination activating gene 1) in T cells under WT (right column) and *Bcl11b* KO (left column) conditions.

**Supplementary Figure 9:**
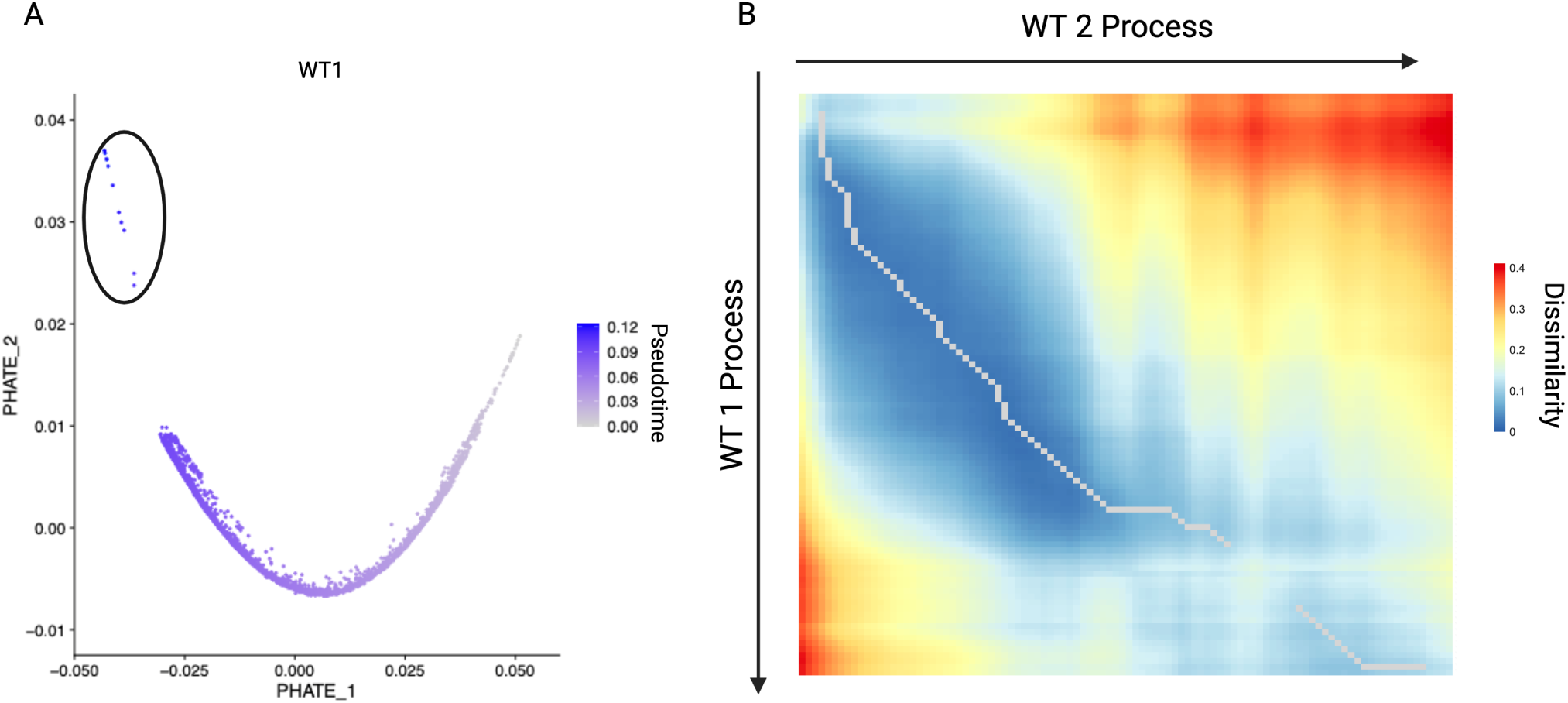
Impact of outlier cells on TrAGEDy alignments. Plot showing the PHATE embeddings of the WT1 T cell dataset with each cell coloured by its cell type annotation. Circle identifies cells which are separate from the main body of the trajectory (A). TrAGEDy alignment of the WT1 and WT2 T cell datasets when the cells circled in A are kept in the trajectory. Dissimilarity in gene expression of interpolated points was calculated using Spearman correlation with blue meaning low dissimilarity and red meaning high dissimilarity.

**Supplementary Figure 10:**
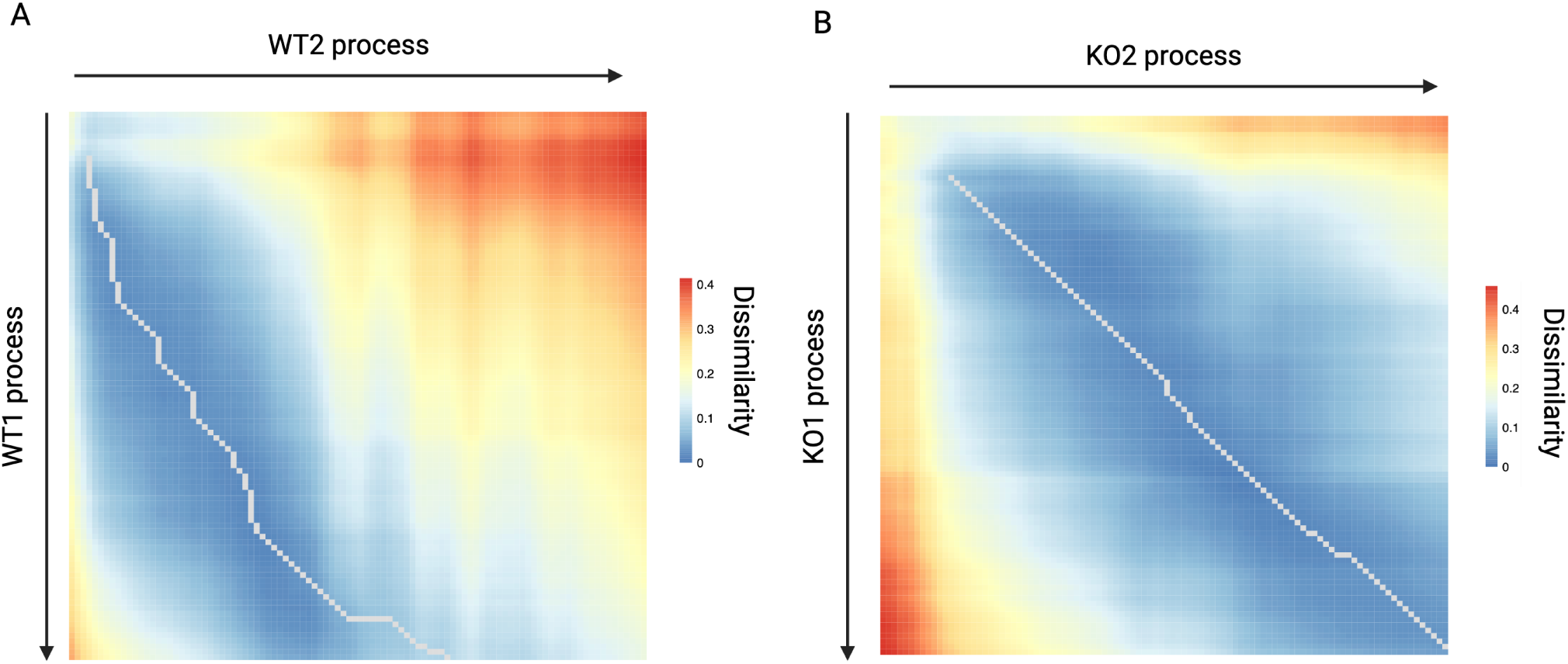
TrAGEDy alignment of WT and *Bcl11b* KO T cell sequencing run trajectories. TrAGEDy alignment of the WT1 and WT2 T cell development trajectories (A) and the *Bcl11b* KO1 and *Bcl11b* KO2 T cell development trajectories (B). Dissimilarity in gene expression of interpolated points was calculated using Spearman correlation with blue meaning low dissimilarity and red meaning high dissimilarity.

**Supplementary Figure 11:**
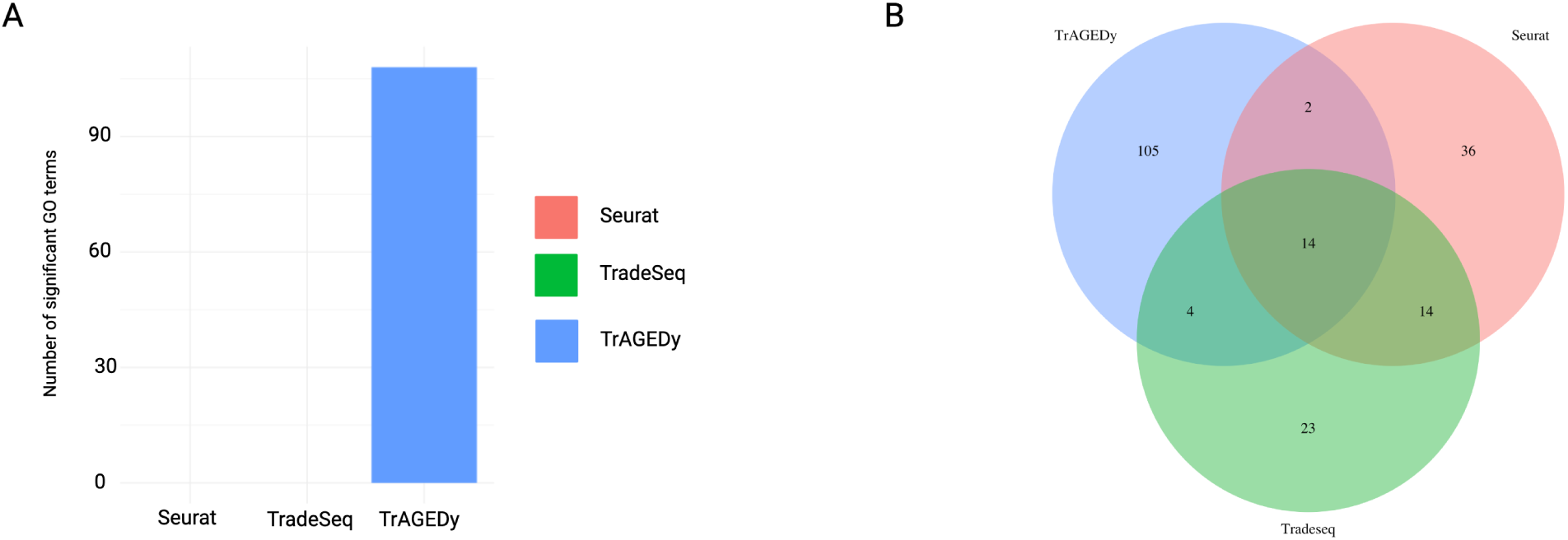
Further GO term analysis of WT vs *Bcl11b* KO T cell development. Barplot showing the number of significant Gene Ontology (GO) terms (Benjamini-Hochberg adjusted p-value < 0.01) returned when GO enrichment analysis was carried out using clusterprofiler on the unique DE genes returned by TrAGEDy, TradeSeq and Seurat (A). Venn diagram showing the intersections of significant GO terms (Benjamini-Hochberg adjusted p-value < 0.01) found by the three methods using clusterprofiler on all the DE genes returned by TrAGEDy, TradeSeq and Seurat (B).

## Notes

### Competing Interest Statement

The authors have declared no competing interest.

### Summary of Updates

Revised abstract to remove the supplementary materials section as it implied we'd been accepted by Bioinformatics, when in fact we've only submitted there.

